# A skyline birth-death process for inferring the population size from a reconstructed tree with occurrences

**DOI:** 10.1101/2020.10.27.356758

**Authors:** Jérémy Andréoletti, Antoine Zwaans, Rachel C. M. Warnock, Gabriel Aguirre-Fernández, Joëlle Barido-Sottani, Ankit Gupta, Tanja Stadler, Marc Manceau

## Abstract

Phylodynamic models generally aim at jointly inferring phylogenetic relationships, model parameters, and more recently, population size through time for clades of interest, based on molecular sequence data. In the fields of epidemiology and macroevolution these models can be used to estimate, respectively, the past number of infected individuals (prevalence) or the past number of species (paleodiversity) through time. Recent years have seen the development of “total-evidence” analyses, which combine molecular and morphological data from extant and past sampled individuals in a unified Bayesian inference framework. Even sampled individuals characterized only by their sampling time, i.e. lacking morphological and molecular data, which we call *occurrences*, provide invaluable information to reconstruct past population sizes.

Here, we present new methodological developments around the Fossilized Birth-Death Process enabling us to (i) efficiently incorporate occurrence data while remaining computationally tractable and scalable; (ii) consider piecewise-constant birth, death and sampling rates; and (iii) reconstruct past population sizes, with or without knowledge of the underlying tree. We implement our method in the RevBayes software environment, enabling its use along with a large set of models of molecular and morphological evolution, and validate the inference workflow using simulations under a wide range of conditions.

We finally illustrate our new implementation using two empirical datasets stemming from the fields of epidemiology and macroevolution. In epidemiology, we apply our model to the Covid-19 outbreak on the Diamond Princess ship. We infer the total prevalence throughout the outbreak, by taking into account jointly the case count record (occurrences) along with viral sequences for a fraction of infected individuals. In macroevolution, we present an empirical case study of cetaceans. We infer the diversity trajectory using molecular and morphological data from extant taxa, morphological data from fossils, as well as numerous fossil occurrences. Our case studies highlight that the advances we present allow us to further bridge the gap between between epidemiology and pathogen genomics, as well as paleontology and molecular phylogenetics.

## 1 Introduction

Birth-death processes are stochastic processes used to model population dynamics with two main parameters, the birth rate and the death rate, which are respectively the rate at which new individuals appear, and the rate at which individuals are removed from the process. In macroevolution, these two rates correspond to the speciation and extinction rates, while in epidemiology they correspond to the transmission and recovery rates. These processes already enjoy a long history of applications in evolutionary biology. In the first half of the twentieth century, Yule (1925) introduces them in the field with macroevolutionary applications in mind, to model the number of species within genera. Kendall (1948) then derives analytically the transition probabilities for linear birth-death processes, and discusses their use in the context of evolutionary biology, with a special focus on epidemiology. Ground-breaking work by Nee et al. (1994) followed on the probability density of the *reconstructed tree* in a linear birth-death process, i.e. the tree obtained by pruning all extinct lineages from the full genealogical history of the process (see Fig. 1A). The linear birth-death process was then later extended to allow rates to vary in different parts of the tree (Alfaro et al. 2009), over time (Morlon et al. 2011), or depending on some character of interest (Maddison et al. 2007).

**Figure 1:**
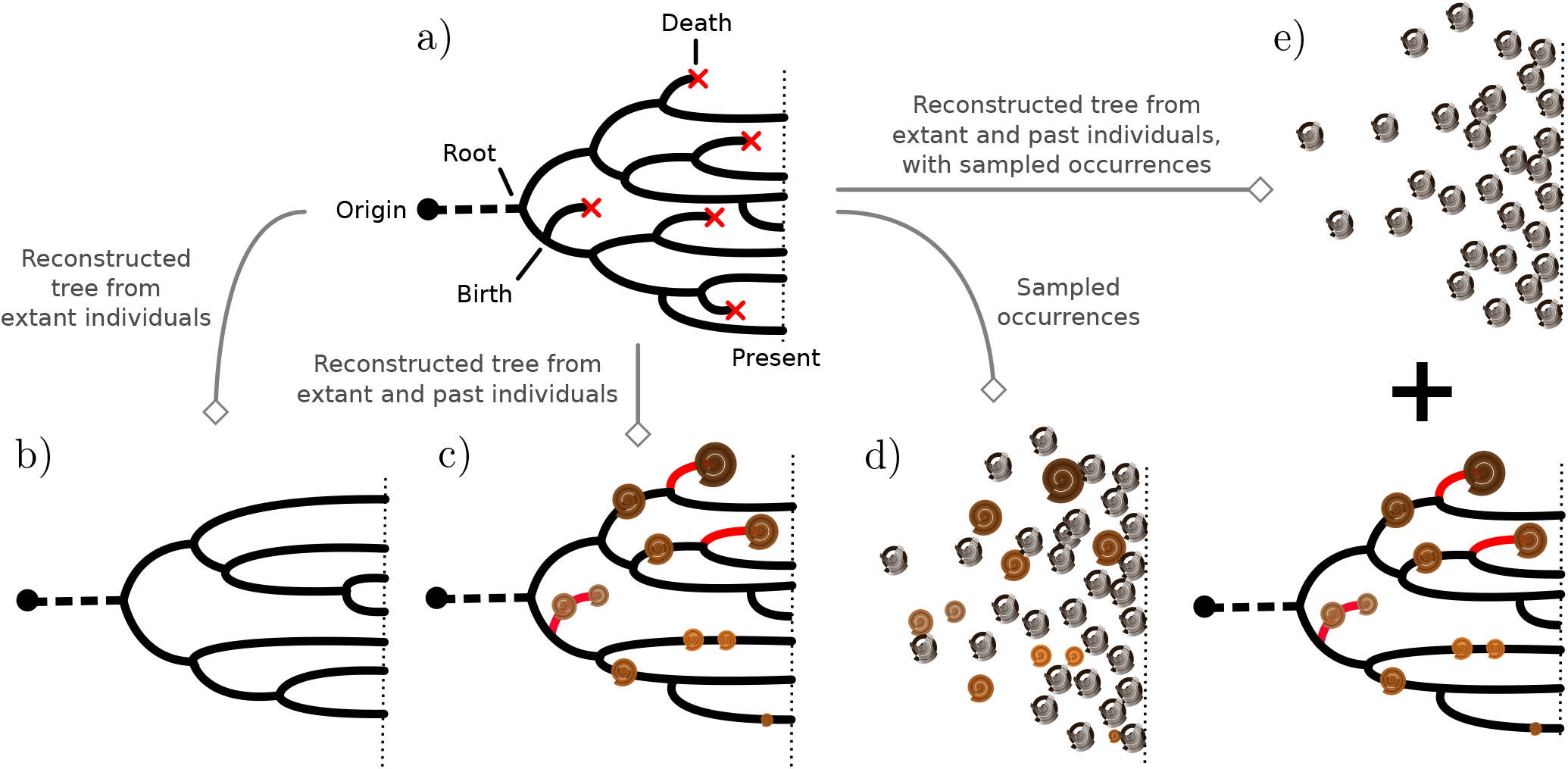
Different approaches to infer past history and population sizes. a) The full unknown history of the population. Different types of data can be used in order to infer the past history and population sizes of the population. b) Genetic sequencing data and character data for present day individuals allows to infer the reconstructed phylogenetic tree and to estimate the past number of lineages. c) This tree can be enriched with past individuals with documented character data (either morphological or molecular data), adding information on some extinct lineages (red). d) Past population sized can be obtained from sampled occurrences alone. Finally, e) A more comprehensive total evidence method integrates extant genetic sequences, samples with character data and the occurrences in a unified framework.

Although diversification histories inferred from extant species sometimes agree with those inferred from the fossil record (Morlon et al. 2011; Xing et al. 2014; Silvestro et al. 2018), there largely remains a gap between these two approaches in macroevolution (Marshall 2017). On the one hand, extant species provide invaluable information regarding the dynamics of the diversification process, especially close to the present. On the other hand, the fossil record, albeit scarce, could much better inform extinction estimates (Quental and Marshall 2010). An extension introduced by Stadler (2010) and dubbed the *Fossilized Birth-Death Process* (FBD) (Heath et al. 2014) has allowed us to model jointly extant and extinct taxa along the same tree, and thus helped bridge the gap between paleontology and molecular phylogenetics. In this model, each species can be sampled throughout its lifetime at a fixed rate, and appear in the reconstructed tree (see Fig. 1B). The probability density of the resulting phylogeny is derived in closed-form, and has been successfully used as a prior in Bayesian phylodynamic analyses to study the diversification history of hymenopterans (Zhang et al. 2015), as well as the penguins (Gavryushkina et al. 2016). The same model was also used in the context of epidemiology, where infected individuals can as well be sampled throughout the infectious period and appear in the reconstructed tree Stadler et al. (2013). Finally, model extensions have been introduced to help take into account the age of species (i.e. stratigraphic ranges) or, in the context of epidemiology, an extended period of infection (Stadler et al. 2018).

An important feature of many standard paleontological datasets, is that only a fraction of fossils have been thoroughly described and are associated with morphological data. Similarly, in standard epidemiological surveys, only a fraction of the recorded case count data is typically sequenced. In this paper, we call *samples with character data* the subset of samples with either morphological data or molecular data, and *occurrences* the recorded samples without character data. This data, while providing no useful information regarding the topology of the tree, still contain invaluable information regarding the underlying population size (see Fig. 1C). For this reason, they have long been used in paleontology to infer diversity trajectories (Raup 1972; Sepkoski et al. 1981), and even preservation, origination and extinction rates in an alternative Bayesian setting (Silvestro et al. 2014, 2019). Some authors have analyzed occurrences in the standard FBD-based Bayesian framework, considering them as leaves in the tree with missing character data, and integrating over the unknown topology (Heath et al. 2014; Gavryushkina et al. 2014; O’Reilly and Donoghue 2020). However, in the event both samples with character data and occurrence data are available, applying the standard FBD model requires making the assumption that both sets of data were generated under the same process and with the same rate. A second step towards integrating these occurrences was performed by Vaughan et al. (2019), who explicitly modeled an additional sampling process for occurrences, allowing for the joint analysis of the observation of a phylogeny and a record of occurrences (see Fig. 1D). Vaughan et al. (2019) additionally proposed an inference framework based on the use of a particle filter to compute the likelihood. Rasmussen et al. (2011) presents another method based on a particle filtering algorithm to consider occurrences and trees in tandem, although in a coalescent framework instead of a birth-death framework. Gupta et al. (2019) built on previous work by Vaughan et al. (2019) and described soon after a fast algorithm to compute the likelihood of the data, focusing on a special case of the model where all individuals sampled through time are removed from the process upon sampling. Finally, under the same model assumptions, Manceau et al. (2019) presented a method to compute the population size distribution conditioned on a reconstructed tree and a record of occurrences.

In this paper, we extend these last two methods to include piecewise-constant parameters, allowing us to explicitly incorporate known variation in birth, death and sampling rates through time. We implement our work as a new distribution, coined the Occurrence Birth Death Process (OBDP), available in the Bayesian phylogenetic software RevBayes (Höhna et al. 2016) to compute the joint probability density of a tree and a record of occurrences. This can readily be used to sample the posterior of trees and population sizes through time, given an observed record of occurrences and a list of samples with character data attached. We illustrate the versatility of the method on two empirical datasets coming from the fields of epidemiology and macroevolution. In epidemiology, we infer the prevalence through time for the Covid-19 outbreak on the Diamond Princess cruise ship, based on the joint observation of molecular sequences and case count data. In macroevolution, we infer the diversity through time in the Cetacean clade, based on the joint observation of molecular data for extant species, morphological character data for some fossils and some extant species, and the record of fossil occurrences available on the Paleobiology Database.

## 2 Material and methods

### 2.1 Phylodynamic model

We consider that a population of individuals starts at the time of origin *t*_or_ with one individual, and evolves through time under a birth-death process with piecewise constant birth rate, λ_*t*_, and death rate, *μ_t_*. Three different sampling schemes are successively applied along the process. First, individuals can be sampled through time and be included in the tree, with piecewise-constant sampling rate *ψ_t_*. Second, they can be sampled through time as raw *occurrences* not included in the tree, with piecewise-constant sampling rate *ω_t_*. Third, individuals reaching present time are included in the tree with a fixed probability *ρ*. Finally, upon sampling, individuals are removed with a piecewise-constant probability of removal *r_t_*.

As a result of these three sampling steps, we observe a reconstructed tree 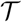, which is the tree spanning all *ψ*-sampled and *ρ*-sampled individuals, as well as a record of occurrences 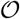, which is a timeline recording successive *ω*-sampling events. We aim at (i) computing the probability density of 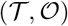, which will play the role of the phylodynamic likelihood in our Bayesian framework, and (ii) compute the probability distribution of the total number of lineages in the process at time *t, I_t_*, conditioned on the observed 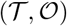. Note that the number of lineages in 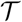 at time *t*, denoted *k_t_*, is an obvious lower-bound of the total number of individuals in the process at time *t, I_t_*. For this reason, we are targeting the probability distribution 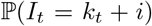, where *i* stands for the number of hidden individuals.

In Appendix A, we extend the method introduced by Manceau et al. (2019) to include piecewise-constant parameters in computing two quantities. First, defining 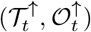 as the tree and record of occurrences constrained to [*t, t*_or_], we aim at numerically computing the joint probability of the partial tree and occurrence record between time *t* and the origin, and the total number of lineages at time *t*,

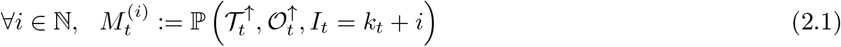

which can be used to compute, upon reaching present day *t* = 0, 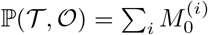.

Second, defining 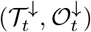 as the tree and record of occurrences constrained to [0, *t*], we aim at numerically computing the probability of the partial tree and occurrence record between time *t* and the present, conditioned on the total number of lineages at time *t*,

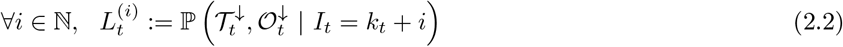

which can as well be used to compute, upon reaching the time of origin *t*_or_, 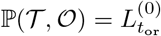.

We derive initializing conditions and Master equations governing the evolution of *M_t_* and *L_t_* through time and compute these quantities by numerically evaluating the system ordinary differential equations (Appendix A). Note that in this numerical evaluation, we have to make one approximation namely, assume a maximal population size *N* (while in theory the population size may become arbitrarily large). In practice, *N* must be chosen large enough to cover most of the high-density support of the *L_t_* and *M_t_* probability distributions to avoid biasing calculations. Finally, provided we know both quantities at time *t*, the probability distribution *K_t_* of the total number of individuals living at time *t* is given by,

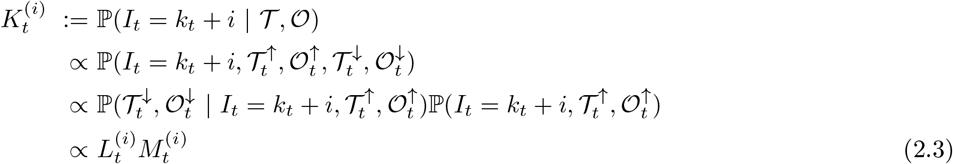

where the last equality is due to the Markov property of the process. We summarize all the notation introduced above in Table 1.

**Table 1:** Parameters and objects of the Occurrence Birth-Death Process.

| Parameter | Signification | Object | Signification |
| --- | --- | --- | --- |
| $t_{or}$ | Time of origin | $\mathcal{T}$ | Reconstructed tree |
| $\lambda$ | Speciation rate | $\mathcal{O}$ | Record of occurrence times |
| $\mu$ | Extinction rate | $I_t$ | Total number of lineages |
| $\psi$ | Fossil sampling rate | $k_t$ | Number of sampled lineages |
| $\omega$ | Occurrence sampling rate | $i$ | Number of hidden lineages |
| $r$ | Removal probability at sampling | $(\mathcal{O}_t^\uparrow, \mathcal{T}_t^\uparrow)$ | Occurrences and tree before time $t$ |
| $\rho$ | Sampling probability at present | $(\mathcal{O}_t^\downarrow, \mathcal{T}_t^\downarrow)$ | Occurrences and subtrees after time $t$ |

### 2.2 Bayesian inference framework

We consider a Bayesian inference framework with additional model layers for character data evolution along the reconstructed tree 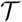. For epidemiology applications, we superimpose a model of molecular evolution, leading to the observation of a sequence alignment for both extant and extinct taxa in 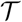. For macroevolution applications, we superimpose (i) a model of morphological evolution, leading to the observation of character data for both extant and extinct taxa in 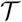, and (ii) a model of molecular evolution, leading to the observation of a sequence alignment for extant taxa only. Summarizing all (molecular and morphological) character data together as 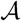, and all model parameters as *θ*, the target posterior distribution of reconstructed trees 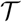 and model parameters *θ* can be written as the product of the phylodynamic likelihood, the likelihood of character data given 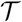 and *θ*, and prior probabilities:

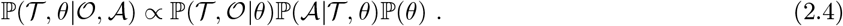

First, we sample this posterior distribution using a Metropolis-Hastings MCMC. Second, the posterior probability distribution of the ancestral population size can be written as,

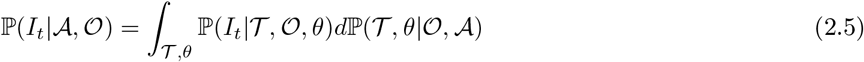

and is thus numerically computed as the arithmetic mean of *K_t_* over the trace of the posterior of 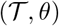.

### 2.3 Numerical implementation

We implement our model in RevBayes (Höhna et al. 2016, 2017), an open-source software for Bayesian inference in phylogenetics. RevBayes is fully based on graphical models (Höhna et al. 2014), a unified framework for representing complex probabilistic models in the form of graphs where nodes correspond to model variables and edges of their probabilistic relationships. It allows the user to construct interactively their own phylogenetic graphical model in the Rev language, by combining the hundreds of available models of nucleotide substitution, rate variation across sites and along the tree, and tree priors proposed in the literature (see Supp Fig. S6B for an illustration with our model). Our three key additions consist of (i) introducing the OBDP distribution (Supp Fig S6A) into RevBayes, so that everyone can use it with their own graphical models; (ii) implementing the core algorithms responsible for computing the quantities *L_t_* and *M_t_* through time and eventually the final log-likelihood; and (iii) including a function to generate the posterior probability distribution of the ancestral population size through time.

Figure 2 summarizes the full workflow to go from the raw data 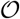, 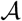 to the inferred reconstructed tree 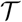, model parameters *θ* and diversity trajectories *I_t_*.

**Figure 2:**
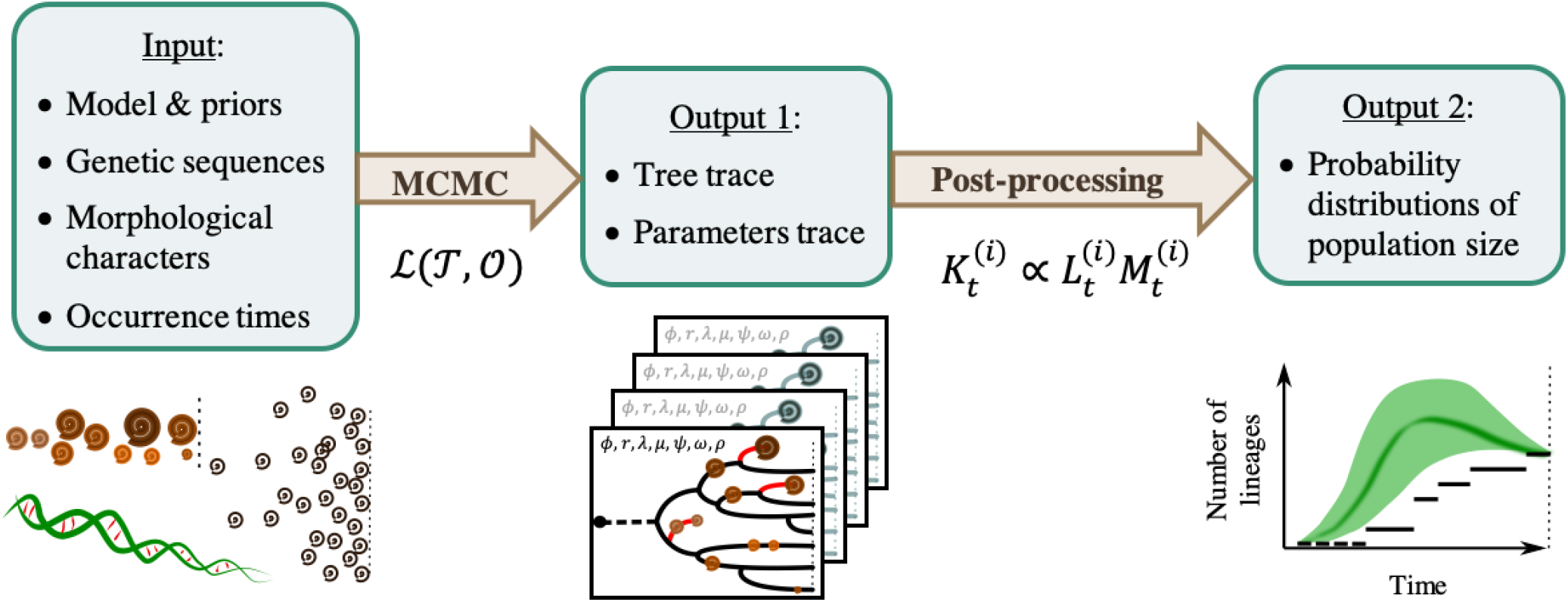
Workflow for using the OBD model for diversity inference. One first needs to specify a graphical model with priors and provide some empirical (molecular, morphological, occurrence) data. A MCMC is run to sample the posterior distribution of trees and parameters. Finally, these traces are used to compute the posterior distribution of diversity through time.

### 2.4 Validation of the method

#### 2.4.1 Direct likelihood comparison

We verify that the phylodynamic likelihood computed using *L_t_* or *M_t_* coincides with (i) previous RevBayes implementations (Höhna et al. 2017; Heath et al. 2019) of linear birth-death processes that are special cases of our framework, when no occurrences are included and *r* = *ω* = 0, and (ii) an earlier Python implementation of the likelihood with constant parameters (Manceau et al. 2019). We use a small fixed dataset and compute the likelihood using (i), (ii) and our implementation, under a wide range of parameters which are listed in Figure 3.

**Figure 3:**
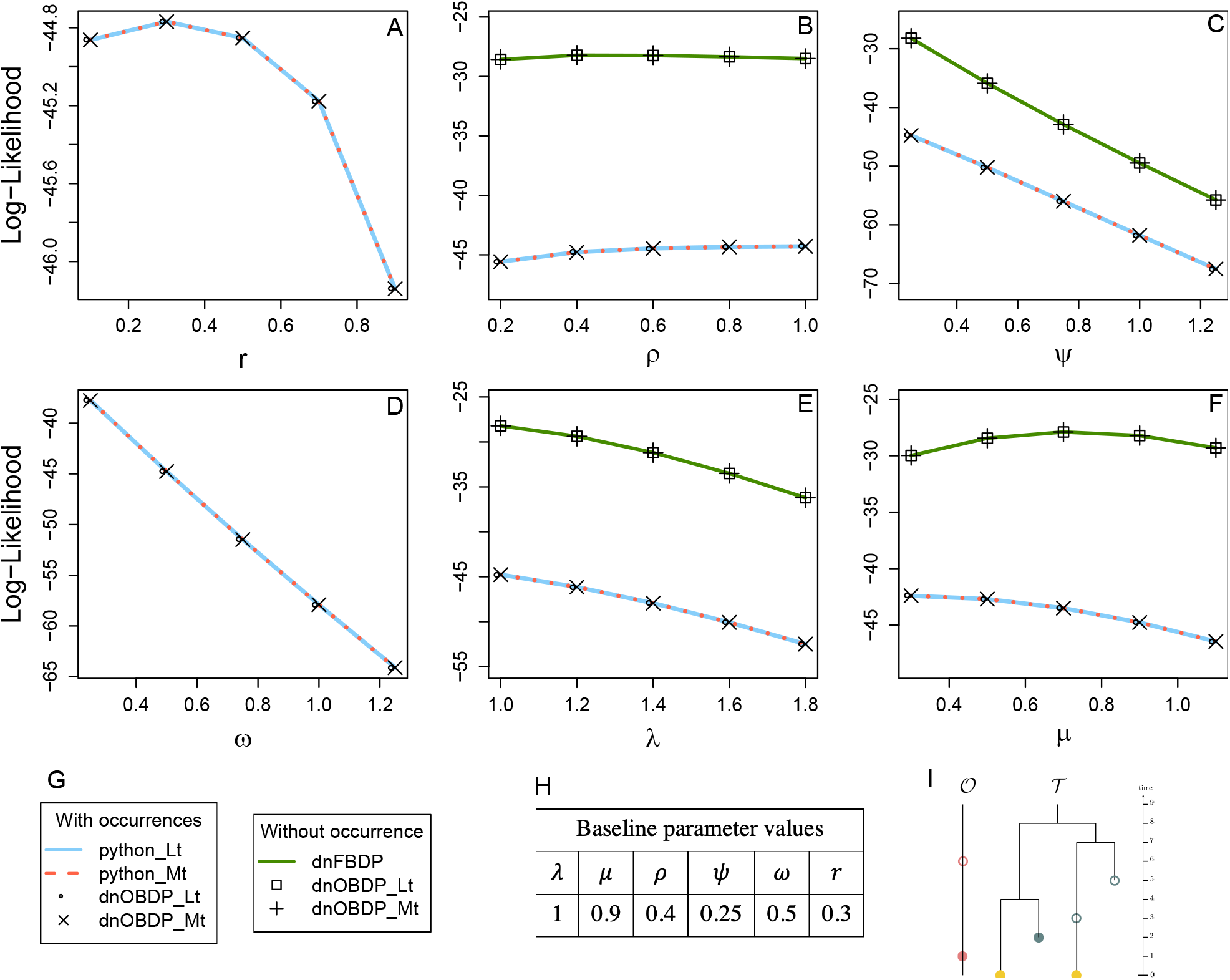
Validation of the likelihood calculation. Each parameter is varying (A-F) while keeping the others at their baseline values (H) and evaluating the likelihood of the toy dataset (I) where pink dots are occurrences, blue dots at past samples, and yellow dots are extant samples. Filled dots are removed samples, unfilled are not removed. For all parameters (A-F), our RevBayes implementation is compared to the Python code provided in Manceau et al. (2019) and whenever possible (B,C,E,F), to earlier FBDP implementations available in RevBayes, fixing *r* = 0, *ω* = 0 and 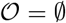.

#### 2.4.2 Qualitative validation on simulated datasets

Time-forward simulations of two full OBD processes were performed to produce two datasets. On the first one (dataset 1), we simulated morphological data for past samples, and both morphological and molecular data for extant samples, mimicking a macroevolution scenario. On the second one (dataset 2), we simulated only molecular data for all samples. Parameter values and the full model specifications are described in Supp Mat C. We used the RevBayes implementation to infer back the parameter values along with the reconstructed tree and population sizes. We performed this validation as a blind test, with two of us being in charge of conducting the analysis, and remaining ignorant of the true simulated scenarios.

#### 2.4.3 Quantitative validation of the MCMC implementation

We follow a procedure called Simulation-Based Calibration (Talts et al. 2018) for validating our MCMC implementation. It consists in the following three steps: (i) we define priors (Table 2) for all the involved parameters and simulate 1000 parameter sets, trees with sampled fossils, occurrences and genetic sequences (100 nucleotides long); (ii) for each simulated dataset, we use the same priors to infer the posterior distribution of reconstructed trees and parameters; and (iii) we compute the proportion of datasets for which the true (simulated) parameter values fall within a 100*α*% credible interval of the posterior distribution, for a range of *α* values (19 evenly spaced between 0.05 and 0.95). If the MCMC is correctly sampling the posterior distribution, the proportion of posterior credible intervals recovering the truth should be close to *α*.

**Table 2:**
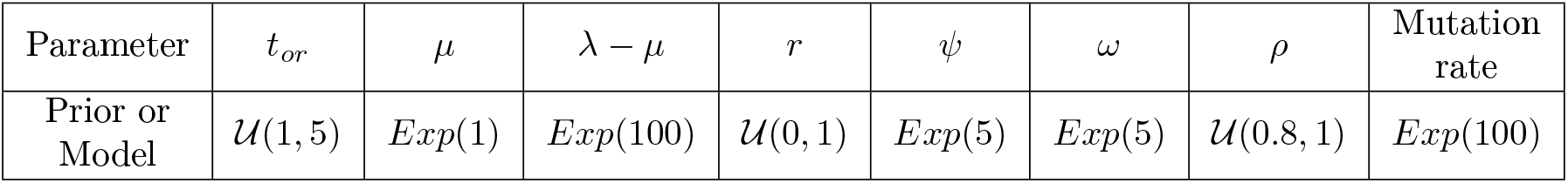
Prior distributions of the OBDP parameters for the quantitative validation test. Notations: *U* for Uniform distribution with given lower and upper bound, *Exp* for Exponential with given rate parameter. The model of molecular evolution is the Jukes-Cantor 1969 substitution model (JC69) with strict clock hypothesis.

| Parameter | $t_{or}$ | $\mu$ | $\lambda - \mu$ | $r$ | $\psi$ | $\omega$ | $\rho$ | Mutation rate |
| --- | --- | --- | --- | --- | --- | --- | --- | --- |
| Prior or Model | $U(1, 5)$ | $Exp(1)$ | $Exp(100)$ | $U(0, 1)$ | $Exp(5)$ | $Exp(5)$ | $U(0.8, 1)$ | $Exp(100)$ |

### 2.5 Covid data analysis

#### 2.5.1 Molecular and occurrence dataset

We use the model implementation with piecewise constant rates to perform a phylodynamic analysis of the SARS-CoV-2 epidemic aboard the Diamond Princess cruise ship, a well-documented outbreak from February 2020. The outbreak is an example of a closely monitored, geographically constrained closed population, and thus constitutes an ideal case study of the disease dynamics and the mitigation policies undertaken (Mallapaty 2020).

The sequenced data used for this analysis consists of a set of 71 full length viral genomes collected between February 15th and February 17th, all acquired from GISAID (Shu and McCauley 2017). Acknowledgements for laboratories that contributed the genome sequences used in this analysis are given in Supp. Mat. E.1. All available sequences were aligned to reference genome MN908947 and sites subject to low sequencing accuracy were masked. Following the standard NextStrain (Hadfield et al. 2018) pipeline, sites 13402, 24389 and 24390 as well as 150 bases at the ends of the genomes were masked, thought to be sequencing artefacts that would bias the alignment.

In this example, we define occurrences as patients testing positive for SARS-CoV-2 using polymerase chain-reaction (PCR) detection methods, recorded as case counts. These case counts, i.e. the daily reports of new cases, along with the total number of samples tested were published by the Japanese Ministry of Work throughout the outbreak; case counts were then compiled in the JHU CSSE database (Dong et al. 2020). Out of all 712 cases detected amongst passengers, we focus on the 705 cases detected while guests were still aboard the cruise ship, from the beginning of the cruise on January 20th until February 27th. Sequencing dates and case counts were communicated as daily reports throughout the outbreak. For all report entries, exact dates were randomly assigned to all occurrences within each day. Additionally, we shift dates by a day to account for the delay between sampling and reporting of the PCR results. The full dataset, in its original and processed formats, is presented in Figure S12.

#### 2.5.2 Model assumptions

The model parametrisation allows us to examine two complementary aspects of the temporal change in epidemic spread. First, we estimate the effective reproductive number across all time intervals of interest. The reproductive number is the expected number of secondary cases produced by a single infected individual and is a standard epidemiological parameter, quantified in our model as 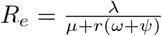. Second, we infer the corresponding prevalence trajectories, aiming to gain insight into the number of potentially undetected patients that are thought to make up a significant proportion of the infected population (Mizumoto et al. 2020).

To achieve these two goals, we make full use of the skyline implementation of our model by allowing independent shifts for each rate parameter. In doing so, we closely follow the exact timeline of events of the outbreak. We first incorporate information regarding the testing and sequencing procedures. The testing strategy was initiated by Japanese authorities after a first guest was confirmed positive for SARS-CoV-2 on February 3rd. It was then extended to asymptomatic passengers from February 11th onward. Sequencing of some of the viral samples was then performed between February 15th and February 17th. To these 4 sampling parameters shifts, we additionally introduce another shift for the birth rate λ before the start of mandatory cabin isolation, on February 5th, producing the full timeline of *m* = 5 intervals.

Reports of the total number of samples tested were assembled to adjust prior means for *ω* + *ψ* on different time intervals, and account for the extension of testing to asymptotic passengers. In total, testing efforts yielded 4066 samples over the entire period of interest, with 3622 of them being obtained after February 11th. All other settings and priors used in this analysis are presented in detail in Supp. Table S7.

### 2.6 Cetacean data analysis

#### 2.6.1 Context

Cetaceans are a group of marine mammals, represented by 89 living species, that possess a remarkable and well-studied fossil record (Fordyce 2009). Their history can be summarized by three main phases (Marx et al. 2016), (i) starting 53 Ma, a 10 Myr land-to-sea transition accompanied by drastic morphological transformations in the archaeocetes (stem cetaceans), (ii) the emergence of neocetes (crown cetaceans, including filter-feeding mysticetes and echolocating odontocetes) at the Eocene-Oligocene boundary (34 Ma) and their radiation up to a Mid-Miocene peak (12 Ma) followed by (iii) a sharp decline in diversity in the last 4-6 Ma.

Several studies have already attempted to estimate the diversity trajectory of cetaceans, using the fossil record (Uhen and Pyenson 2007), molecular phylogenies (Morlon et al. 2011) or both (Marx and Fordyce 2015); but even the latter total-evidence study did not include all fossil occurrences in its analyses. The initial huge discrepancies between the history inferred from the fossil record and from molecular phylogenies (Quental and Marshall 2010) have been partially bridged, but including occurrences may help provide a more reliable time-calibrated tree and a robust diversity trajectory estimation.

#### 2.6.2 Molecular, morphological and occurrence datasets

The data can be subdivided in three parts: molecular, morphological, and occurrences. Datasets were collected and analysed separately and are stored on the Open Science Framework (https://osf.io) (Aguirre-Fernández et al. 2020). Molecular data comes from Steeman et al. (2009), and comprises 6 mitochondrial and 9 nuclear genes, for 87 of the 89 accepted extant cetacean species. Morphological data was obtained from Churchill et al. (2018), the most recent version of a widely-used dataset first produced by Geisler and Sanders (2003). After merging 2 taxa that are now considered synonyms on the Paleobiology Database (PBDB) and removing 3 outgroups that would have violated our model’s assumptions, it now contains 327 variable morphological characters for 27 extant and 90 fossil taxa (mostly identified at the species level but 21 remain undescribed). In order to speed up the analysis we further excluded the undescribed specimens and reduced this dataset to the generic level by selecting the most complete specimen in each genera. Indeed, the computing cost increases quadratically with the maximum number of hidden lineages N, to the point of becoming the bottleneck in our MCMC when *N* > 100. Given that a mid-Miocene peak diversity between 100 and 220 species is expected (Quental and Marshall 2010), with less than 100 observed lineages in our inferred tree at that time, N should therefore be about 150. Inferring instead the tree of cetacean genera allows us to reduce *N* to 70 hidden lineages. The final dataset thus contains 41 extant and 62 extinct genera.

Occurrences come from the PBDB (data archive 9, M. D. Uhen) on May 11, 2020. The dataset initially consisted of all 4678 cetacean occurrences, but the cetacean fossil record is known to be subject to several biases (Uhen and Pyenson 2007; Marx et al. 2016; Dominici et al. 2020). A detailed exploration (see Supp. Mat. D) of this occurrence dataset revealed several notable biases. First, an artefactual cluster of occurrences in very recent times, combined with other expected Pleistocene biases (Dominici et al. 2020), led us to remove all Late Pleistocene and Holocene occurrences. Second, we detected substantial variations in fossil recovery per time unit across lineages (see Supp. Fig. S10) resulting from oversampling of some species and localities, possibly due to greater abundance or spatiotemporal biases (Dominici et al. 2020). This observation violates our assumption of identical fossil sampling rates among taxa during a given interval. In order to reduce this bias, we retained occurrences identified at the genus level and further aggregated all occurrences belonging to an identical genus found at the same geological formation. Occurrences for which the geological formation was not specified, we used geoplate data combined with stratigraphic interval. This resulted in a total of 968 occurrences retained for the analysis.

#### 2.6.3 Model assumptions

Each fossil comes along with a stratigraphic age uncertainty interval. Reducing this interval to either the midpoint, or a uniformly drawn point, has been shown to lead to serious biases in the divergence time estimates (Barido-Sottani et al. 2019). We instead follow the same procedure as Heath et al. (2019) and apply a uniform prior for the age of fossils with morphological characters, within the bounds of their stratigraphic age uncertainty. As a result, the age of a fossil included in the tree can slide within this interval during the MCMC.

Based on previous work showing huge discrepancies in mutation rates between odontocetes and mysticetes (Dornburg et al. 2012), and generally between nuclear and mitochondrial sequences (Allio et al. 2017) we partitioned between the two types of sequences and considered a relaxed clock across the tree.

Much less biological knowledge is available about the dynamics of morphological characters (Wright 2019). We thus chose a minimal substitution model and partitioned the alignment in order to treat separately characters that are represented by a different number of states.

All prior distributions are fully detailed in Supp. Table S6.

## 3 Results

### 3.1 Validation of the method

#### 3.1.1 Direct Likelihood Comparison

We illustrate in Figure 3 the perfect agreement with likelihood values computed using previous functions under a wide range of parameters, for both *L_t_* and *M_t_* traversal algorithms.

#### 3.1.2 Qualitative Validation on simulated datasets

On Figure 4, we superimpose the true, simulated, trajectory of the total number of individuals, together with the inferred posterior distribution. Most importantly, the true trajectory falls within, or is very close to the boundaries, of the 95% posterior credibility interval at any point in time.

**Figure 4:**
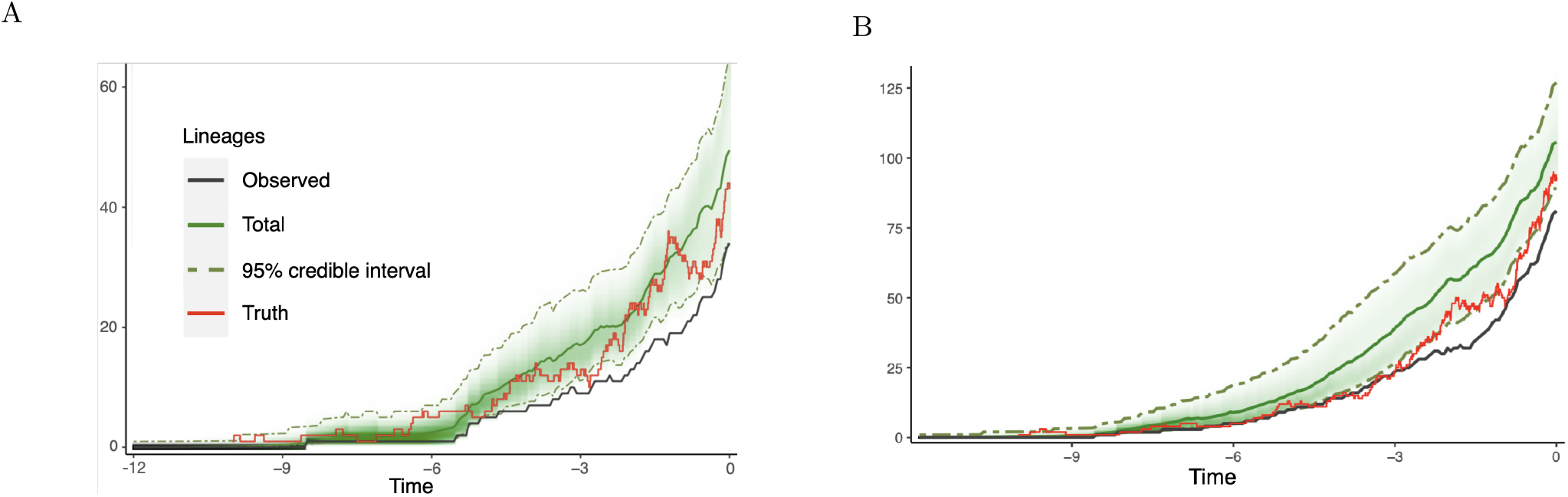
Results of the blind test analysis. The posterior distribution of the number of lineages through time is in green. The inferred LTT plot, showing the number of lineages in the tree through time, is in black. The true simulated number of individuals is in red.

On both datasets, the topology of the tree was also well recovered but divergence dates do not always perfectly match (see Supp. Mat. Figure S9). Due to a greater amount of data in genetic sequences of both past and extant individuals, the divergence dates have been better inferred on dataset 2 as compared to dataset 1.

#### 3.1.3 Quantitative Validation of the Diversity Inference

Figure 5 shows a good correspondence between the proportion of posterior credible intervals containing the true parameter value and the width of the credible interval. This indicates that the MCMC is properly calibrated, i.e. samples adequately the targeted posterior distribution.

**Figure 5:**
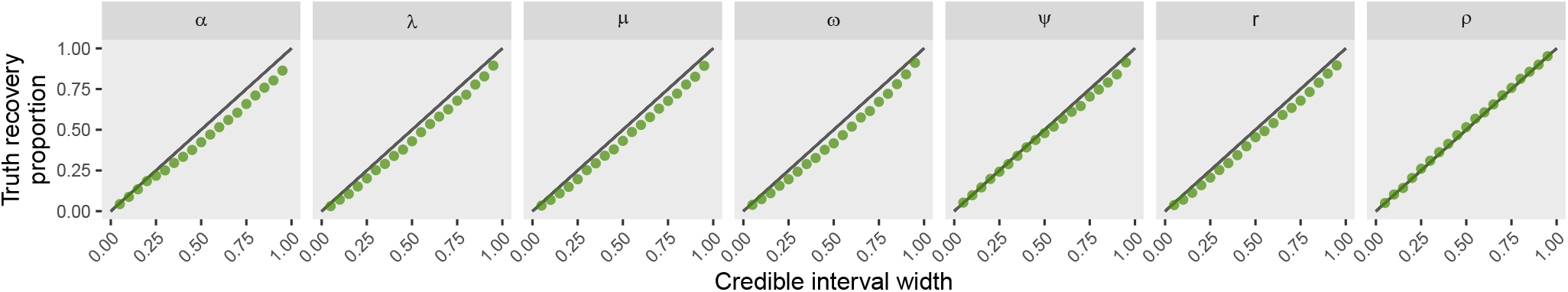
Results of the Simulation-Based Calibration. Each dot corresponds to the proportion of simulated parameters (y axis) falling within its inferred posterior credible interval with a given level (x axis). The black line corresponds to the expected perfect match.

### 3.2 Reproductive number and prevalence in the Covid outbreak

Figure 6 shows the raw data, as well as the estimates of the total prevalence and reproductive number through time. The instantaneous prevalence is typically always slightly lower than the total number of cases sampled each day, which corresponds to the sum of all sampled (and likely removed) individuals over a one-day period. The reproductive number is inferred with very high uncertainty in the beginning of the epidemic, when very few cases were observed, and with a much higher precision in the second part of the process. It decreases synchronously with the launching of non-pharmaceutical interventions in early February (i.e. testing effort and cabin quarantine).

**Figure 6:**
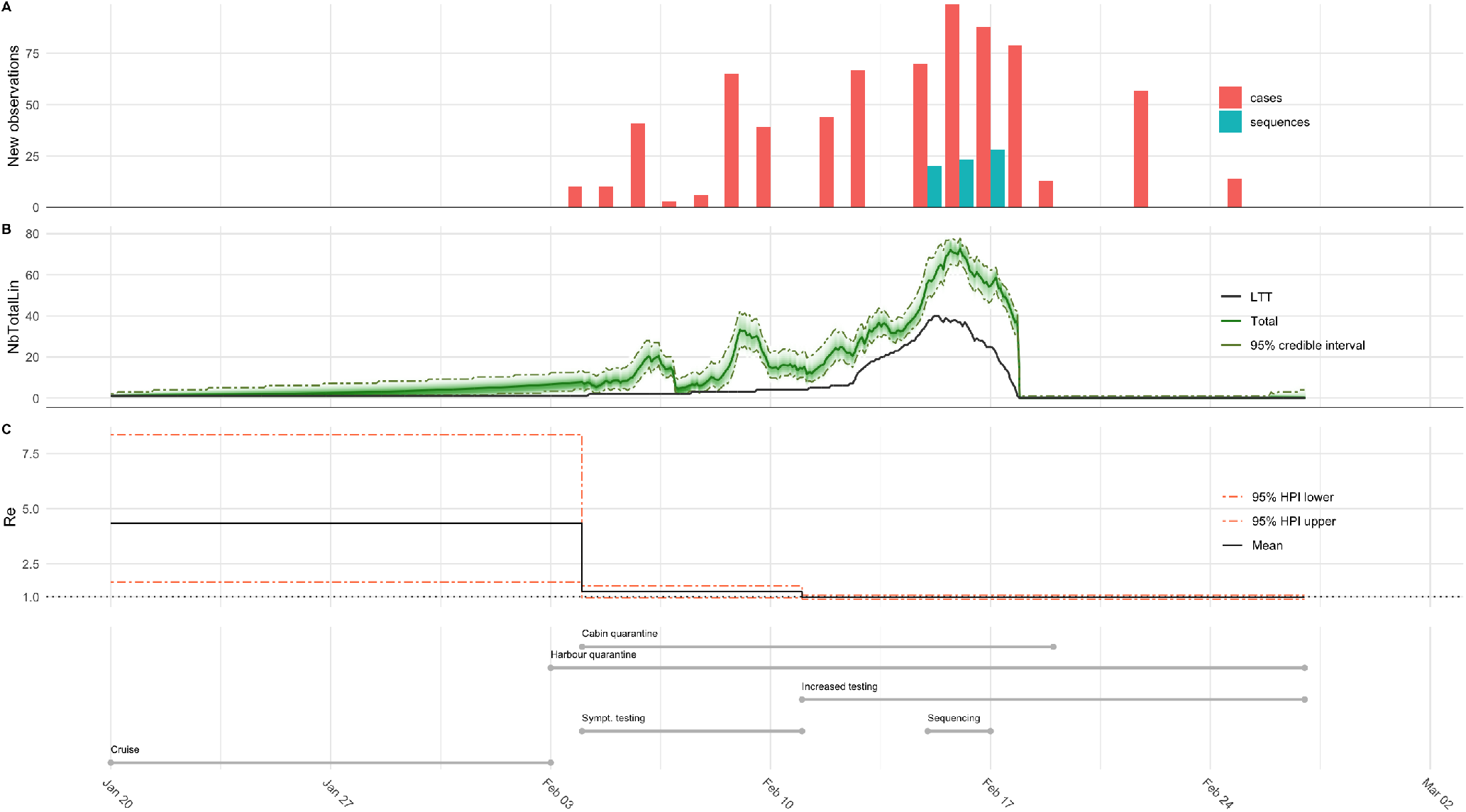
Analysis of the SARS-2 COVID-19 outbreak dynamics aboard the Diamond Princess cruise ship. (A) Occurrence and sequenced data are plotted as daily new observations. Although passengers were monitored until July 2020 (Ministry of Health and Welfare 2020), we focus on infections detected while guests were still aboard, until the end of the harbour quarantine. (B) Posterior probability distribution of the instantaneous total infected population aboard the cruise ship and inferred LTT. (C) Estimates of the effective reproductive number (*R_e_*) throughout a 38 day period starting at the beginning of the cruise.

### 3.3 Total diversity in the Cetacean clade

Figure 7 shows the inferred diversity of cetacean genera over the past 50 Myr. The diversity curve indicates an Early-Eocene origin followed by a monotonous diversification up to a first Mid-Miocene peak (12 Ma), before reaching its maximum in the Pliocene (3.5 Ma) with almost 70 inferred genera. The last million years correspond to a sharp decline leading to the 41 extant genera.

**Figure 7:**
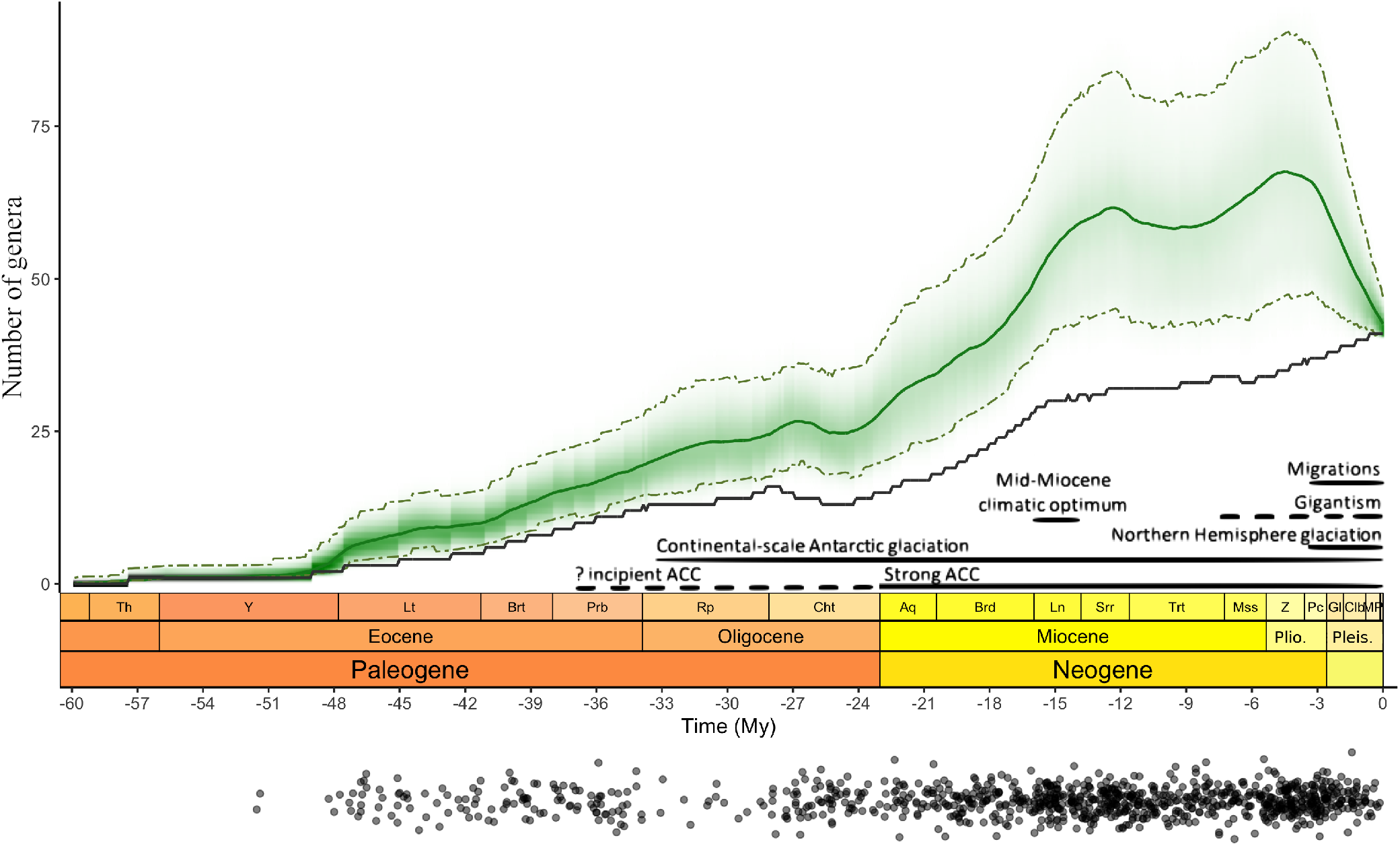
Posterior probability distribution of the total number of genera over time. The 95% credible intervals are indicated in dashed lines, the expected diversity is in green and the inferred LTT is in black. Periods of biotic or abiotic factors that are hypothesized to have driven diversification changes are adapted from Marx and Fordyce (2015) and shown in black for information but had no influence on the analysis. ACC = Antarctic Circumpolar Current. Black dots below represent the occurrences used in the analysis.

## 4 Discussion

### 4.1 Technical achievements and limitations

In this paper, we extend the work of Gupta et al. (2019) and Manceau et al. (2019) to consider piecewise-constant rates through time, and implement the Occurrence Birth-Death Process in the popular phylogenetic inference software RevBayes. This enables us to simultaneously incorporate numerous occurrences without character data, together with taxa for which we have genetic sequences and/or morphological characters. In addition to using the OBDP as a tree prior for inferring epidemiological or macroevolutionary parameters, tree topology and divergence dates, it allows users to compute the posterior probability distribution of the population size through time, in a post-MCMC analysis (see workflow in Fig. 2). We validate the framework, both qualitatively and quantitatively, and illustrate its use in the fields of epidemiology and macroevolution.

The likelihood computation can be very fast when lineages become extinct upon sampling (*r* = 1), relying on the results of Gupta et al. (2019). In practice, this assumption only makes sense for some epidemiological applications, when infected individuals can self-quarantine and be safely assumed to be removed from the process. For macroevolutionary applications, the *r* parameter typically equals zero, and the likelihood computation relies on a more computationally intensive method to numerically solve Master equations (see details in Supp. Mat. A).

More work is thus needed to help speed up the likelihood computation when *r* ≠ 1, on datasets for which a large number of hidden lineages is expected. This will be especially important for further applications in macroevolution and paleobiology, as many data sets feature thousands of fossils occurrences.

### 4.2 Covid-19 Diamond Princess epidemic

The application of our method to the study of a thoroughly documented outbreak highlights the versatility of our model implementation. The ability to incorporate both incidence data and pathogen sequences, in combination with temporal information constitutes one of the first few instances of the use of a *total evidence* approach for the inference of epidemiological trajectories.

In fact, encouraging conclusions can be drawn from both our parameter estimates and the corresponding trajectory inference. First, the reproductive number is inferred to be 4.33 in the absence of any intervention and detection, during the first 15 days of the cruise (see Fig. 6B). These values are consistent with previous studies estimating the basic reproductive number *R*_0_ between 2.5 and 3.5 for the global pandemic (Stadler 2020a), and 3.53 for the Diamond Princess outbreak before quarantine (Stadler 2020b). Second, we infer a decrease in the reproductive number, that remains near or below 1 in the last 23 days of the time period of interest, suggesting that the epidemic was contained by measures taken. Interestingly, we note that this decrease is driven by the sampling and removal of individuals from the infectious population, with the total sampling rate going up from 1.41 to 1.79 days^−1^, after February 11th (see Figure S13 for detailed timeline). This extended sampling, stemming partly from the decision to test asymptomatic passengers, results in occurrences having a visible impact on the reconstructed prevalence curve. This underlines successful integration of both sequence and occurrence data, and the added value brought about by this new implementation.

Biases inherent to many epidemiological data sets indicate important areas for development. For instance, sampling of outbreaks is most often carried out with the aim of quickly monitoring the disease, without rigorously following a protocol. This can result in inconsistent sampling and reporting strategies, with gaps and/or missing data. Due to a 24 hours reporting delay for this dataset, the first detected case was, for example, originally placed after the start of the quarantine. We tried to meticulously remove as many such biases as possible, but our framework could be improved in the future to account for these.

Other potential biases include the effect of population structure – the Diamond princess outbreak is likely to have spread in at least two distinct sub-populations: guests and crew members (Nishiura 2020) – and density dependence – the outbreak being in a closed, geographically constrained population (Rocklöv et al. 2020). Further developments of the method to cover these scenarios could provide even better insight into the dynamics of this outbreak.

### 4.3 Past cetacean diversity

Molecular and paleontological data come with their inherent limitations, e.g. lack of information about extinct lineages for the former and substantial spatiotemporal biases for the latter. Combining them into a single analysis may gather enough signal to mitigate these limitations, but special attention should be paid to model assumptions. We have endeavoured to respect these constraints, by (i) correcting occurrence distribution sampling biases, and (ii) making the most of the piecewise-constant parameter framework to include shifts in diversification rates, as detected previously (Rabosky 2014), as well as shifts in fossilization rates corresponding to the well-established low preservation rates in the Early Oligocene (Rupelian), Early Miocene (Aquitalian) and End Miocene (Messinian) (Marx et al. 2016).

The emerging patterns of cetacean generic diversification in Figure 7 are coherent with previous estimates (Uhen and Pyenson 2007; Morlon et al. 2011; Marx and Fordyce 2015): (i) the “boom and bust” dynamics of prolonged diversification followed by a recent decline is recovered, and (ii) estimated generic richness is higher than the incomplete raw generic counts, as expected. The diversification of cetaceans, starting in the Eocene and accelerating in the Neogene, has been associated by previous authors with the development of the Antarctic Circumpolar Current (ACC) that fuelled a diatom radiation, via nutrient supply, prompting the diversification of bulk filtering cetaceans. The diversity drop in the last 4 million years has been linked to the global climate deterioration and the Northern Hemisphere glaciation, which coincides with the final establishment of modern mysticete gigantism and long-distance migration. Our inferred diversity trajectory (Fig. 7) is compatible with these hypotheses. On the other hand, the distinct second peak with maximum diversity in the Pliocene is unexpected and will require further investigation. Similar to applications in epidemiology, insights into macroevolution based on our novel framework will also benefit from developments that account for biases in fossil sampling, e.g. spatial and temporal biases (Close et al. 2020).

### 4.4 New avenues for phylogenetics

Over the last decade, the field of phylogenetics has expanded considerably with the development of the Fossilized Birth-Death Process and related extensions, of which the Occurrence Birth-Death Process is the latest instance. As a result, the long-standing opposition between molecular-based and fossil-based macroevolutionary inferences is in the process of being bridged, and case count records can be analyzed jointly with sequencing data in epidemiology applications. Many extant clades with a relatively rich paleontological record – e.g. turtles, sharks, angiosperms – as well as outbreak surveillance data, could benefit from this new method to infer reliable phylogenies and diversity/prevalence trajectories.

Future progress could be made to couple birth rates with abiotic drivers, such as biogeography (see also work on multitype birth-death processes Scire et al. (2020)), or biotic drivers such as density-dependence (see also Etienne et al. (2012)). Going even further down the mechanistic road for macroevolutionary applications, stratigraphic palaeobiology could even become an explicit part of diversification models, by considering the accumulation of sediments over finer time and spatial scales (Patzkowsky and Holland 2012). Birth-death process models also exist for the analysis of stratigraphic ranges, or paleontological data only (Stadler et al. 2018; Silvestro et al. 2019), further expanding the potential of phylogenetic models in quantitative paleobiology. We anticipate that these approaches will all benefit from combining paleontological and molecular data.

Overall, our two empirical applications demonstrate that a phylogenetic framework can be successfully applied to recover both the past outbreak prevalence and the past paleodiversity. In contrast to alternative approaches, it maximises the use of available evidence, since it uniquely allows us to combine genetic an morphological character data, together with occurrences. Further, our inference method relies on a generating model, incorporating explicit assumptions about the processes giving rise to our data, including sampling, and is prospectively much more flexible than alternative approaches to mitigating sampling biases.

## Appendix A skyline birth-death process for inferring the population size from a reconstructed tree with occurrences

This appendix presents the detailed derivation of the model used in “A skyline birth-death process for inferring the population size from a reconstructed tree with occurrences” by Andréoletti, Zwaans et al, as well as supplementary results and figures. We extend results of Gupta et al. (2019) and Manceau et al. (2019) to piecewise-constant parameters, describe our implementation in the RevBayes software, and give detailed information on all priors used for simulation or inference in our analyses.

## A Method extension to piecewise-constant parameters

### A.1 Notation and outline of the general strategy

We first recall in Figure S1 the notation that we introduced in the main text with the three different sampling (*ψ*-sampling for sampling of fossils with inclusion in the tree, *ω*-sampling for occurrences and *ρ*-sampling at present).

**Figure S1:**
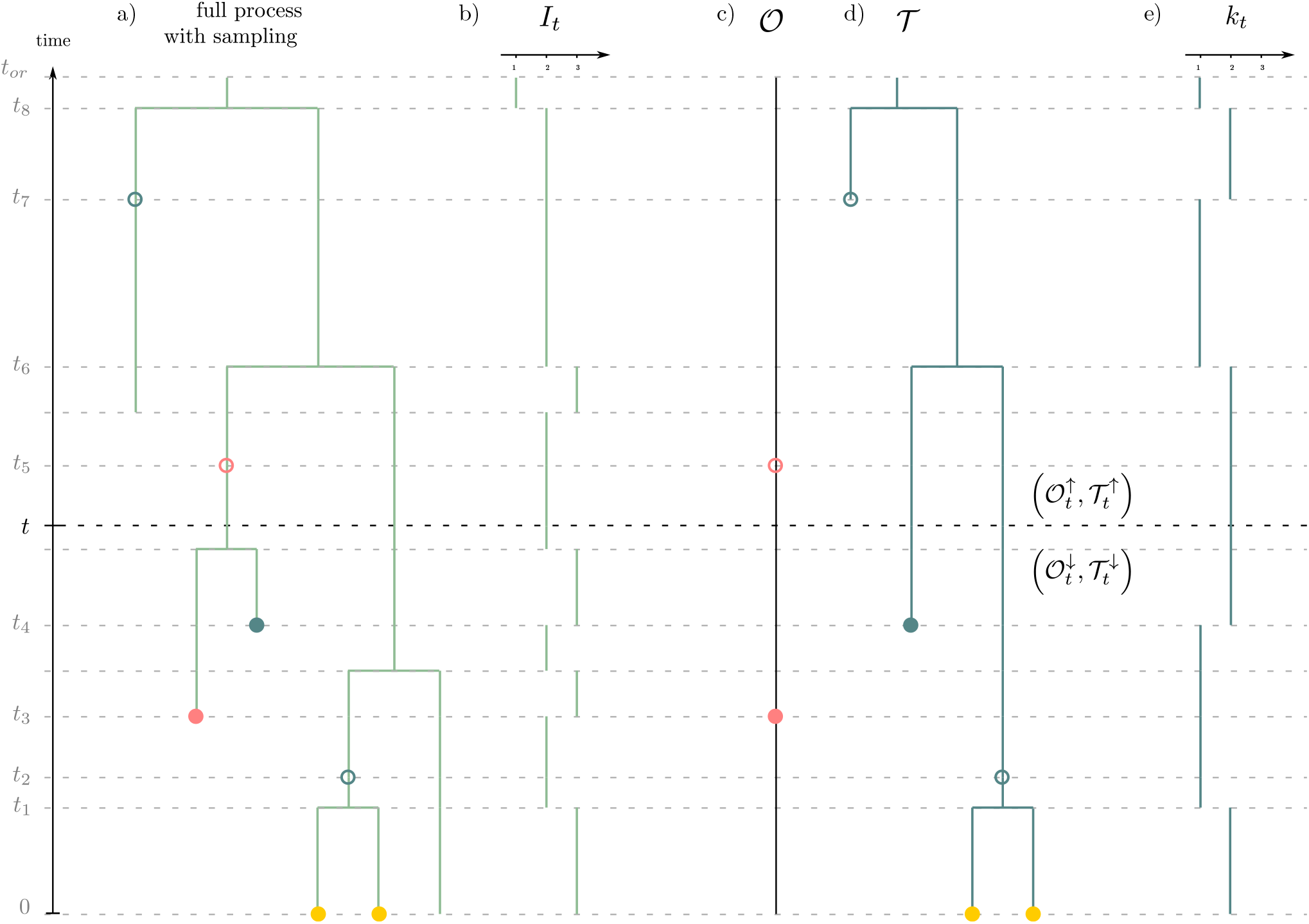
General setting of the method. a) the full process with sampling. Pink dots correspond to ω-sampling (sampling through time without sequencing), blue dots correspond to *ψ*-sampling (sampling through time with sequencing) and yellow dots correspond to *ρ*-sampling at present. Filled or unfilled dots correspond respectively to sampling with or without removal. b) Total number of individuals through time. c) Record of occurrences. d) Reconstructed tree spanning *ψ*- and *ρ*-samples. e) Number of lineages through time in the reconstructed tree (i.e. LTT plot).

To compute the likelihood of 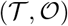 under this process, we will slice horizontally our observations and perform a breadth-first traversal of these. We thus introduce now,

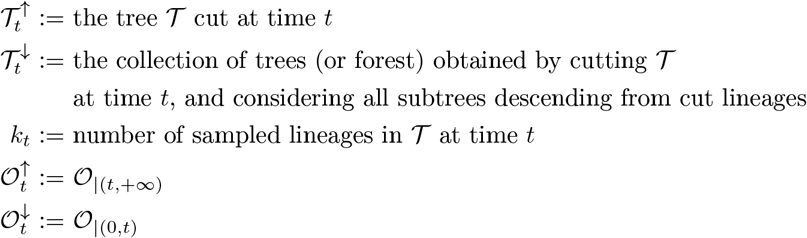

We can now recall the definition of our two key probability densities,

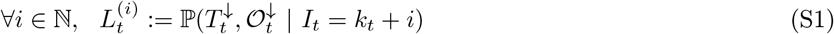

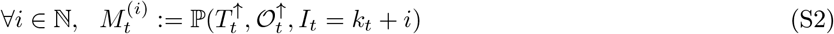

These probability densities have been introduced in Manceau et al. (2019) as a way to target the probability distribution *K_t_* of the total number of individuals given the data. Indeed,

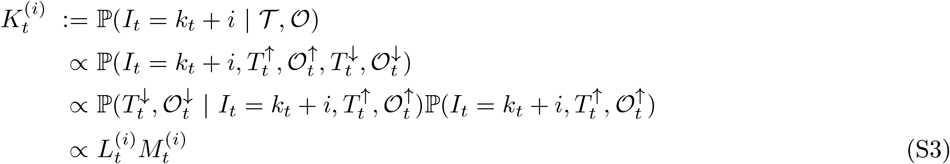

The general strategy of the methods consists of (i) traversing the data backward in time to compute *L_t_*; (ii) traversing the data forward in time to compute *M_t_*; (iii) using the results to compute *K_t_*. This scheme is illustrated in Figure S2.

**Figure S2:**
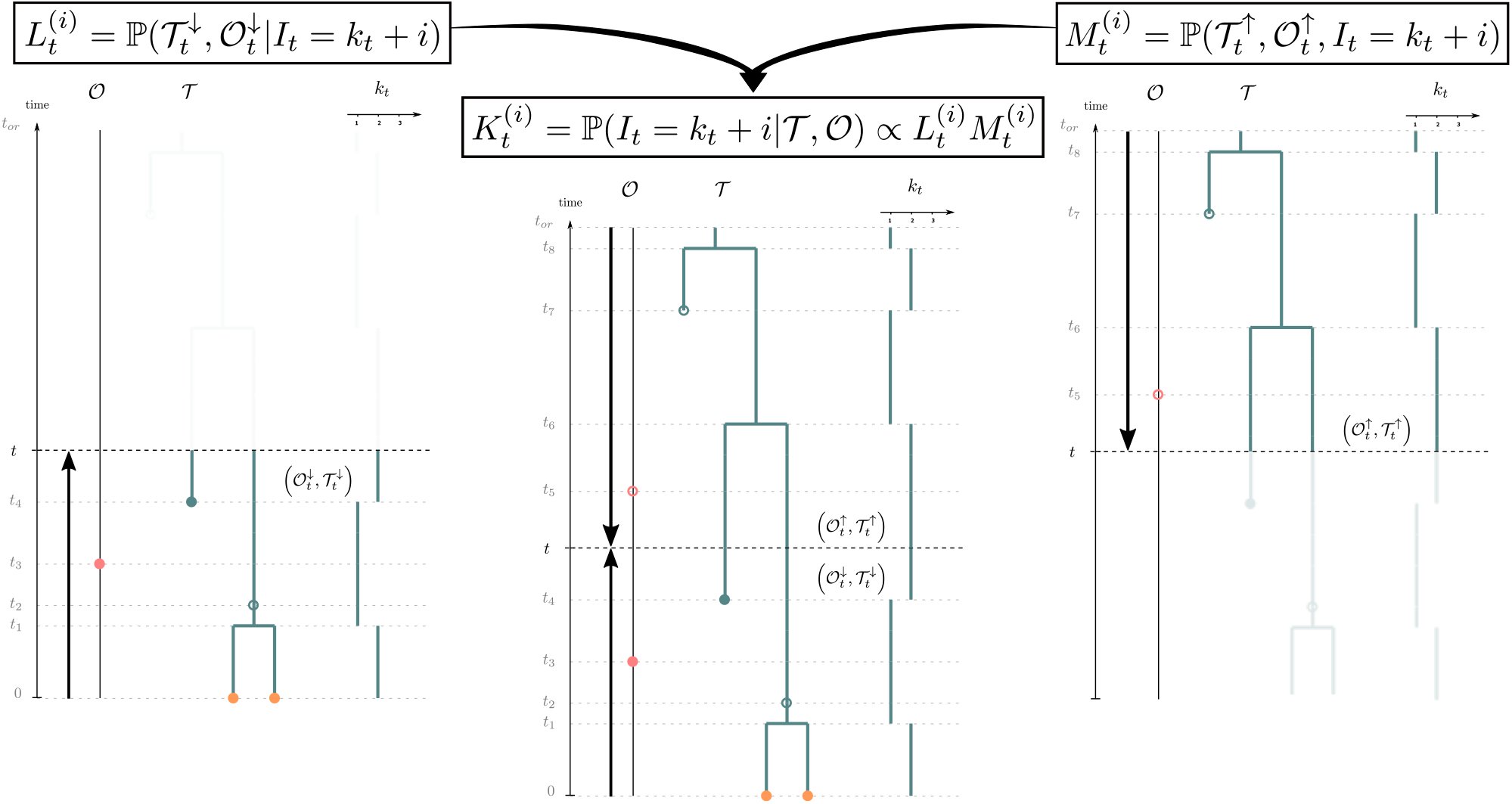
Inferring the posterior distribution of the number of individuals (*K_t_*) in the OBDP. The probability distribution of the past number of lineages *K_t_* is obtained at each time t by combining the quantity *L_t_* obtained from the backward traversal algorithm (left) and the quantity *M_t_* obtained from the forward traversal algorithm (right). See Table 1 for notations.

In the rest of this appendix section, we present the Master equations governing the evolution of these densities through time in a setup with piecewise-constant parameters.

### A.2 Temporal setup for piecewise constant parameters

We partition time into two distinct units.

First, we define periods of time with no observations or sampling events, coined *epochs*, which allow for the basic derivation of Master equations of *L_t_* and *M_t_*. Epochs are delimited by all *n punctual events* times (i.e. branching and sampling events) in 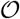 and 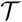 pooled in an ordered list 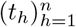. Epoch *h* is thus defined as the time interval (*t_h_, t*_*h*+1_).

Second, we account for all rate shift events, which define *constant rate time intervals*. If we have *m* such intervals, we pool all *m* + 1 rate shift events in an ordered list 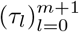, where by convention we consider that *τ*_0_ = 0 and *τ*_*m*+1_ = *t_or_*. Rate time interval *l* is defined as (*τ_l_, τ*_*l*+1_), with parameter set (*λ_l_, μ_l_, ψ_l_, ω_l_, r_l_*). We illustrate this setup in Figure S3 below.

**Figure S3:**
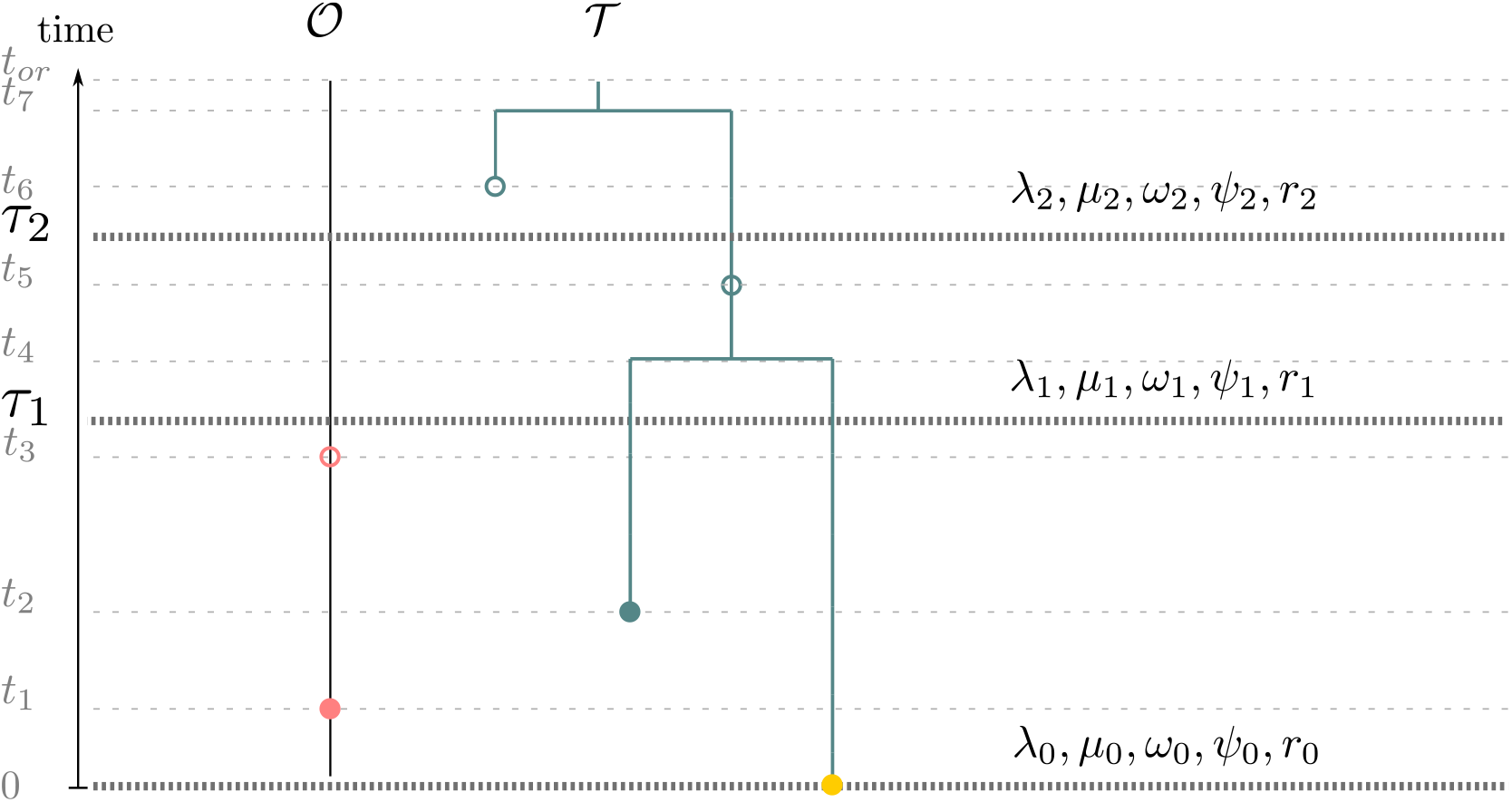
Temporal setup of the method.

### A.3 Master equations governing *L_t_* and *M_t_*

Probability densities *L_t_* and *M_t_* satisfy different Master equations obtained by studying their evolution through time along any given epoch. These are ordinary differential equations (ODE) that can be approximated numerically. Here, we assume *τ_l_* ≤ *t* < *τ*_*l*+1_ meaning that parameters have values (*λ_l_, μ_l_, ψ_l_, ω_l_, r_l_*).

First, we can initialize *L_t_* and *M_t_* respectively at present time 0 and at the time of origin *t_or_*. At present, *ρ* sampling of extant tips yields,

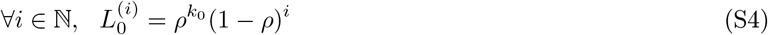

while at the time of origin, the process starts with only one individual *k_tor_* = 1, which yields,

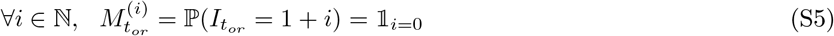

We now consider all events happening in an infinitesimal time step *δt* in the full underlying process which do not result in observations or samplings. Three scenarios correspond to this case:

1. nothing happened with probability (1 – *γ_l_*(*k* + *i*)*δt*), where *γ_l_* = *λ_l_* + *μ_l_* + *ψ_l_* + *ω_l_*
2. a birth event happened:

a. among the *k* sampled lineages in 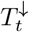, and it leads to an extinct or unsampled subtree to the left or to the right with probability 2*λ_l_kδt*
b. among the *i* other individuals with probability *λ_l_iδt*.
3. a death event happened among the i particles, with probability *μ_l_iδt*

We combine these to write, 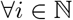,

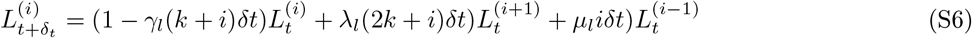

Letting *δt* → 0 yields the following differential equation for *L_t_*,

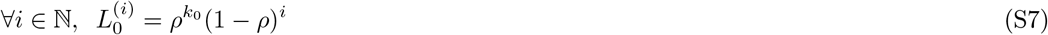

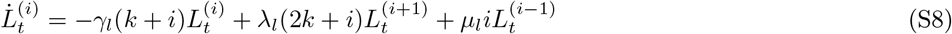

Similarly, *M_t_* is the solution of the following ODE,

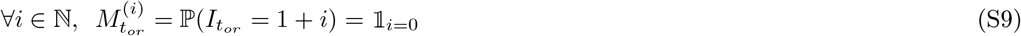

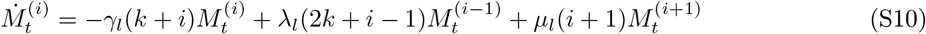

### A.4 Updates at punctual events

**Figure S4:**
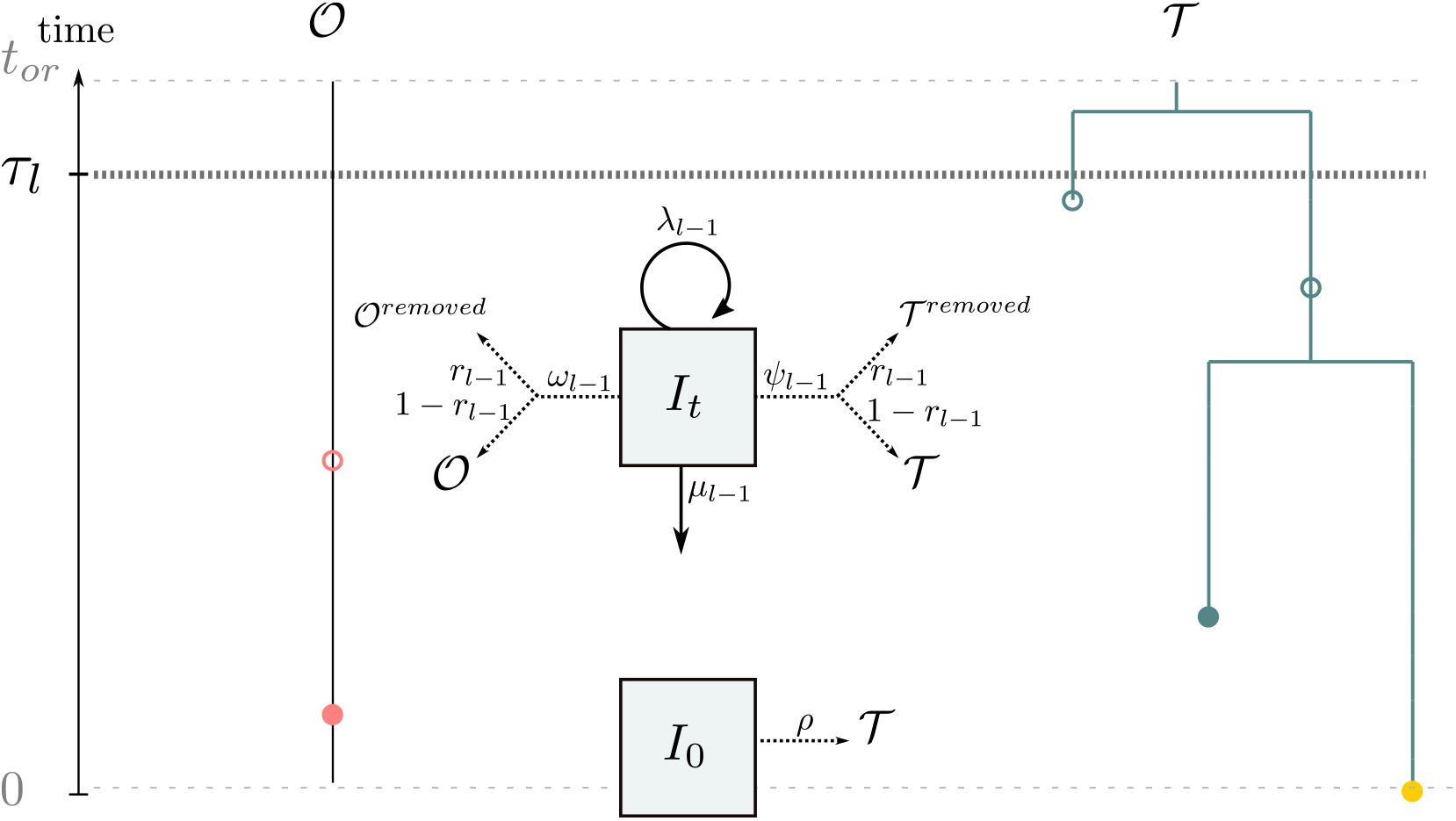
Updated sampling scheme of the method.

There are 6 types of punctual events in 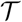 and 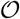 that affect the probability densities *M_t_* and *L_t_*. These correspond to all different sampling options along 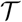 and 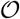 as illustrated in Figure S4. We denote as *M*_t^−^_ and *L_t^−^_* the probability densities immediately prior to the event and *M_t^+^_* and *L_t^+^_* immediately after each event. We emphasise that the expressions differ when considering the process forward in time for *M_t_* or backward in time, for *L_t_*. These cases are the following:

1. sampling of a leaf:

a. in 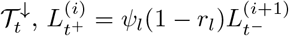
b. in 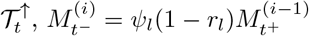
2. removed sampled leaf:

a. in 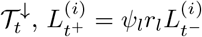
b. in 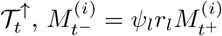
3. sampling along a branch:

a. in 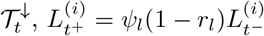
b. in 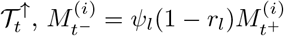
4. occurrence:

a. in 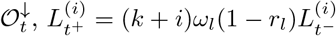
b. in 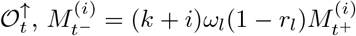
5. removed occurrence:

a. in 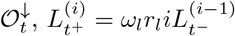
b. in 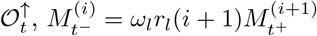
6. branching event:

a. in 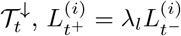
b. in 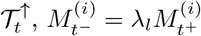

### A.5 Numerical approximation of the ODEs

As described in A.3, for any constant rate time interval where *τ_l_* ≤ *t* < *τ*_*l*+1_, *M_t_* and *L_t_* are defined along epochs as the solution to systems of differential equations S8 and S10 for *t_h_* ≤ *t* < *t*_*h*+1_. Numerically, the solution to such systems of equations is approximated by truncating the system at a fixed integer *N* as follows:

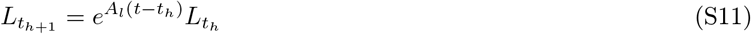

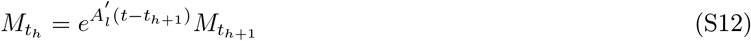

Where *A_l_* and 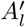 are *N* × *N* tridiagonal matrices with ODE coefficients. When there is a rate shift *τ_l_* within an epoch (*t_h_, t*_*h*+1_), the epoch is cut in two parts and *L_t_* and *M_t_* are simply computed as,

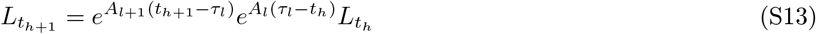

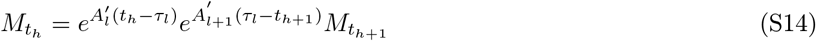

This can be extended to any number of rate changes within an epoch. This strategy of solving for *L_t_* and *M_t_* yields the following two algorithms. Because exponential matrices are computationally intensive to calculate, these algorithms are only used in the most general cases, when no other analytical formula is available (i.e. when *ω* = 0 and *r* = 1).

**Algorithm 1.**
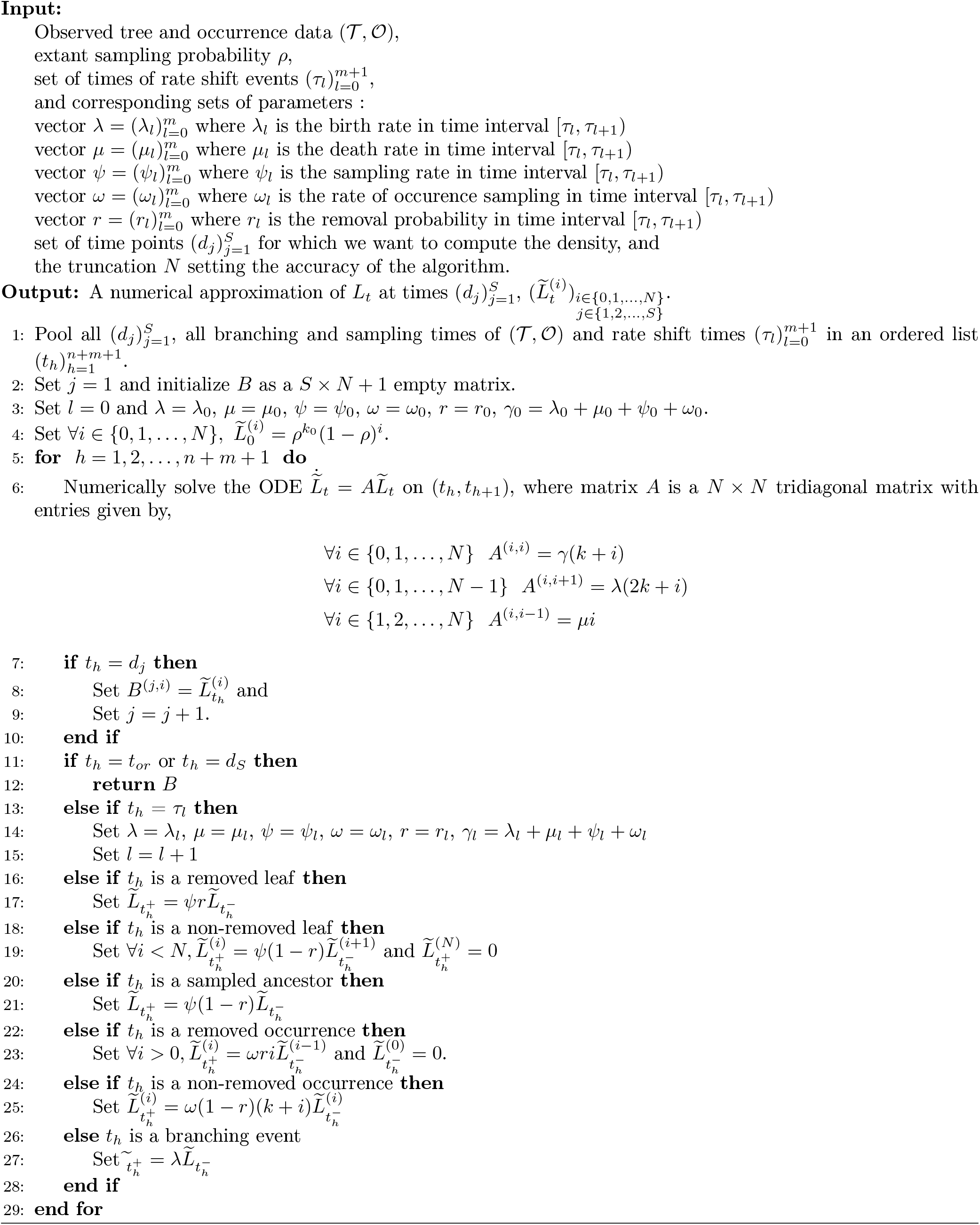
Computes a numerical approximation of *L_t_* for a specific set of times with known rate shift events

**Algorithm 2.**
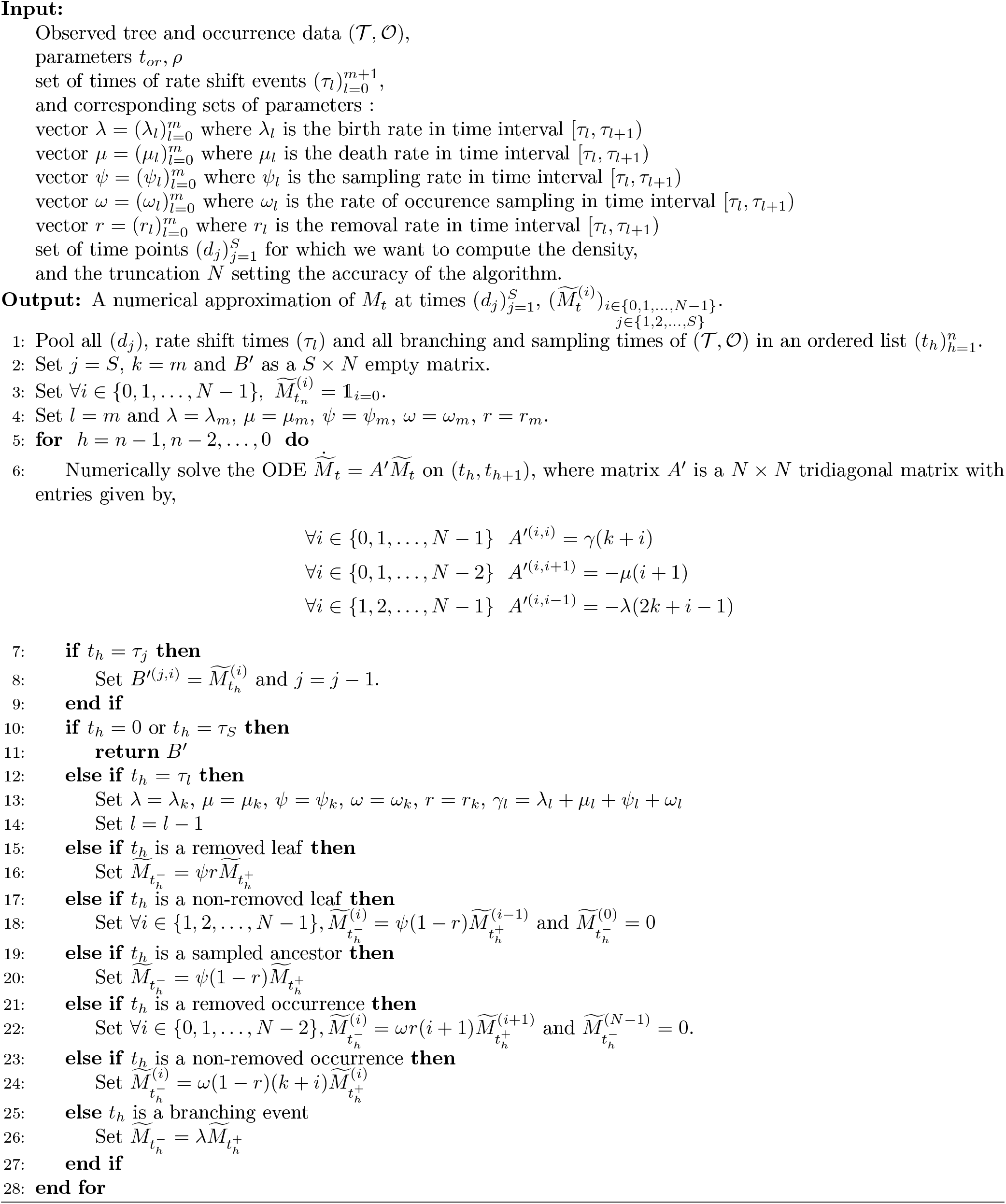
Computes a numerical approximation of *M_t_* for a specific set of times with known rate shift events

## B RevBayes implementation

### B.1 Core algorithms

To enable great flexibility and ensure fast computation, RevBayes is constructed around a mirror structure (Figure 5) in which all the core functions coded in C++ are reflected in the revlanguage section that links with the Rev language interface.

**Figure S5:**
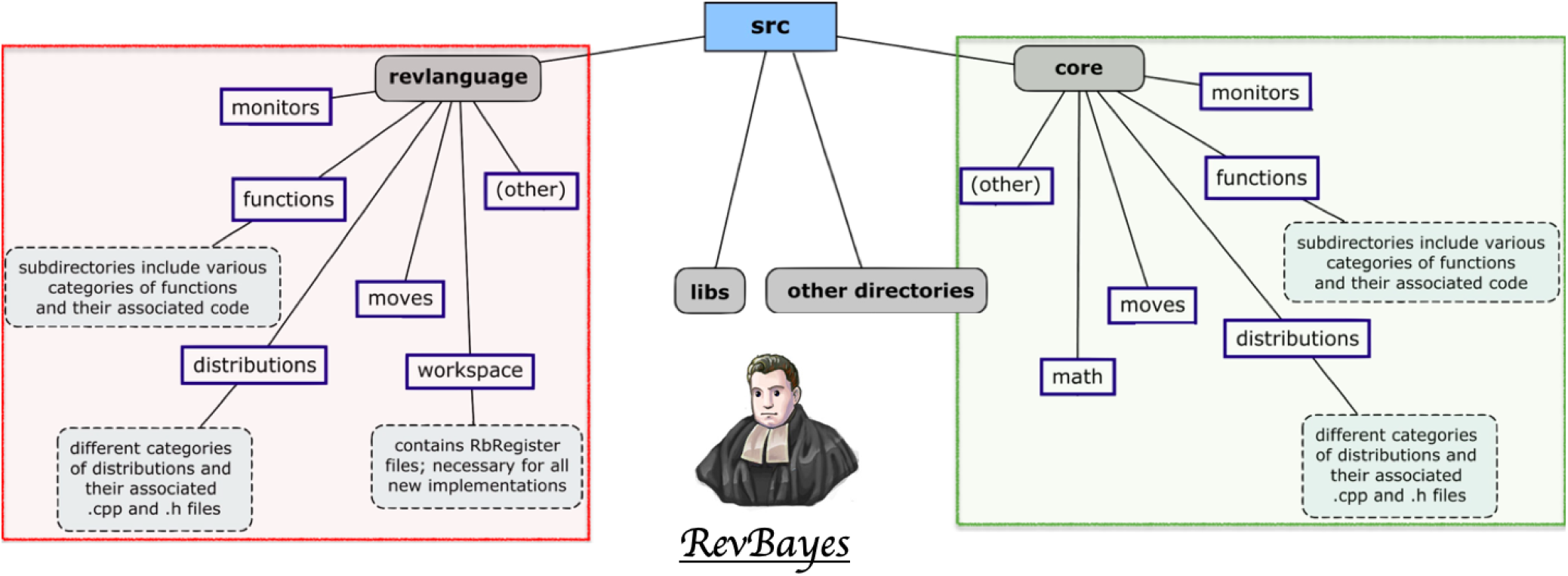
Simplified representation of the RevBayes structure. Modified from the [RevBayes website](https://revbayes.github.io/developer/architecture/), keeping only descriptions of the folders we modified. Note the organizational symmetry between the core directory containing the hard-coded features and the revlanguage directory matching the Rev syntax.

Due the multiple advantages of RevBayes and its increasing use, particularly for macroevolutionary research, we chose this software to implement the OBDP. All our modifications have been carried out in a separate copy of its development branch on GitHub (https://github.com/revbayes/revbayes/tree/dev-cevo-lab), and are aimed to be integrated in a future stable release. They consist in 3 key additions detailed in Table S1.

The necessary first step was to implement the core algorithms responsible for computing the quantities *L_t_* and *M_t_* through time. The final organisation is as follows: from outside of the *ComputeLikelihoodsLtMt.cpp* file (see Supp. Table S1) the only functions called are *ComputeLnProbabilityDensitiesOBDP* – returning *L_t_* and *M_t_* through time – or *ComputeLnLikelihoodOBDP* – returning only the final likelihood. Those functions will themselves call the appropriate internal function (*ForwardsTraversalMt* or *BackwardsTraversalLt*) with the correct parameters. Those rely on a key function, *PoolEvents*, the role of which is to construct the vector containing all the events that will be browsed by the traversal algorithms, namely branching times, *ψ*- and *ω*-sampling times, and time points for which we want to store the probability distribution.

Because the densities computed during the traversals very quickly reached excessively small or elevated values, to the point of exceeding the maximum number of recorded decimals, a correction term is added at each step to bring the densities closer to 1. At the end of the traversal, the recorded correction terms plus the factorizable factors are added to the log-transformed densities.

In addition, the Occurrence Birth-Death Process and the traversal algorithms not only allow us to perform a MCMC phylogenetic inference incorporating the occurrences, they can also be used to output the probability distribution of the population size through time, Kt. We introduced this functionality into RevBayes through *InferAncestralPopSizeFunction*, which can be called directly from the Rev interface. As with the OBDP distribution, we had to design the parameter loading procedure, then call the *ComputeLnProbabilityDensitiesOBDP* function to get the log(*L_t_*) and log(*M_t_*)) matrices and finally combine and normalize them to obtain the log(*K_t_*)) matrix.

**Table S1:** Overview of the implementations carried out to incorporate the Occurrence Birth-Death Process and the associated Diversity Inference method into RevBayes. It lists for each of our goals the associated C++ files, along with their assignment in the RevBayes structure.

| Objectives | Location | File names | Major new functions |
| --- | --- | --- | --- |
| 1. Perform Forwards and Backwards traversal algorithms | core/<br>functions | *ComputeLikelihoods*<br>*LtMt.h*<br>*ComputeLikelihoods*<br>*LtMt.cpp* | *ComputeLnProbabilityDensitiesOBDP*<br>*ComputeLnLikelihoodOBDP*<br>*PoolEvents*<br>*ForwardsTraversalMt*<br>*BackwardsTraversalLt* |
| 2. Encode the OBDP distribution | core/<br>distributions<br>revlanguage/<br>distributions | *OccurrenceBirthDeath*<br>*Process.h*<br>*OccurrenceBirthDeath*<br>*Process.cpp*<br>*Dist_occurrenceBirth*<br>*DeathProcess.cpp*<br>*Dist_occurrenceBirth*<br>*DeathProcess.h* | *OccurrenceBirthDeathProcess*<br>*computeLnProbabilityDivergenceTimes*<br>*createDistribution*<br>*getParameterRules* |
| 3. Infer past diversity | core/<br>distributions<br>revlanguage/<br>distributions | *InferAncestralPop*<br>*SizeFunction.h*<br>*InferAncestralPop*<br>*SizeFunction.cpp*<br>*Func_inferAncestral*<br>*PopSize.h*<br>*Func_inferAncestral*<br>*PopSize.cpp* | *InferAncestralPopSizeFunction*<br>*createFunction*<br>*getArgumentRules* |

**Figure S6:**
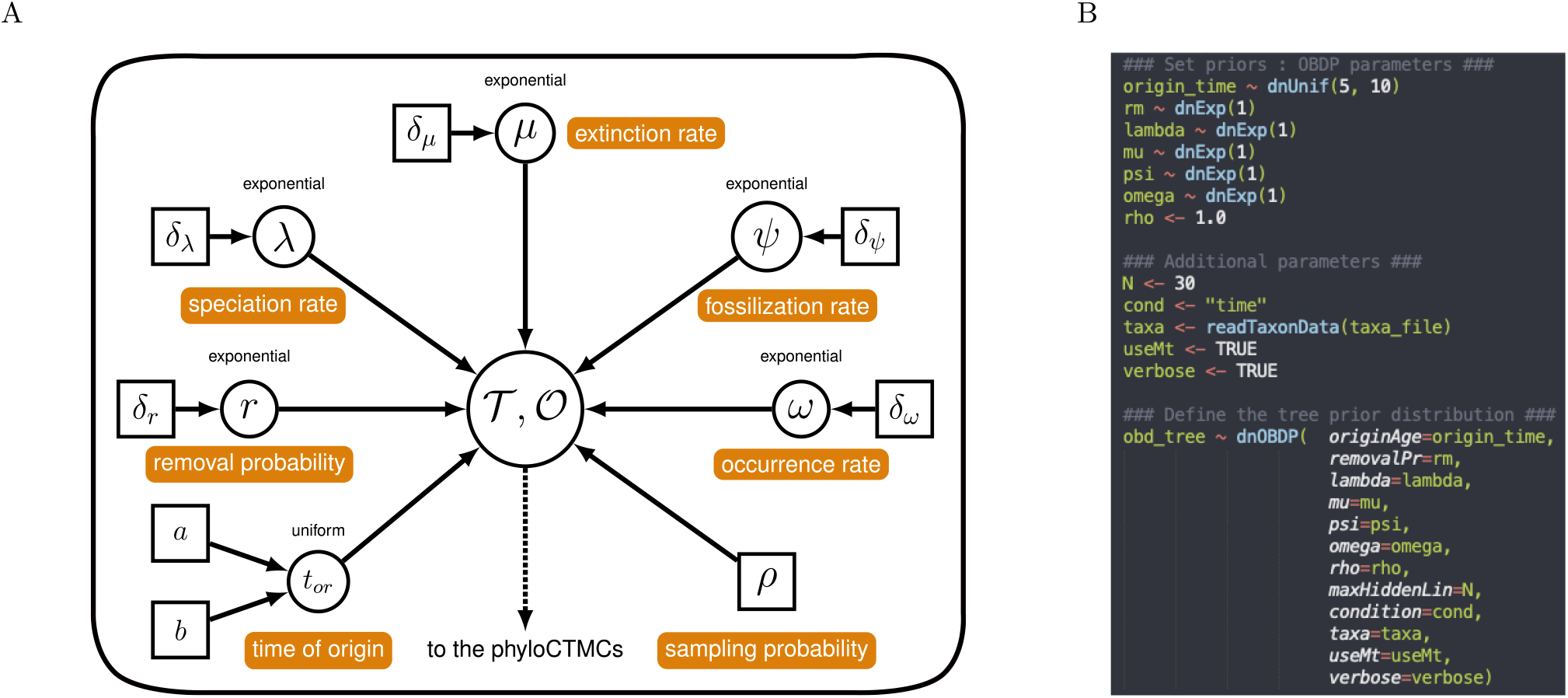
A graphical model of the OBDP and its translation into the Rev language. (A) Graphical model, modified from the RevBayes FBD tutorial, representing the OBDP parameters – labelled in orange – generating a reconstructed tree 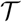 and a record of occurrences 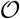. (B) Rev script corresponding to this graphical model. Note the distinction between the ~ notation attributing a distribution to a stochastic node and the ← notation defining a constant node.

### B.2 RevGadgets

The postprocessing step consists in computing the posterior probability of the total number of individuals through time. It can be performed independently of the previous steps, given that one has at least a tree, a set of parameters and optionally occurrence times. It comprises 2 steps, the first one uses the *fnInferAncestralPopSize* function, implemented in RevBayes, to obtain the matrix of diversity densities *K_t_* for each tree in the MCMC trace. Then, in order to convert *K_t_* matrices into a nicely rendered plot we added two functions in the auxiliary R package RevGadgets (https://github.com/revbayes/RevGadgets/tree/dev-plotDiversity). Starting from the trace of posterior trees, parameters, and *K_t_* matrices one first needs to execute the *rev.process.nbLineages* function that will organize the required information into the *Kt_mean* data frame. The goal is to incorporate all the uncertainty concerning the inferred parameter values and tree topologies into the diversity trajectory estimation. Afterwards, this averaged *Kt_mean* is used by the function *rev.plot.nbLineages* to realize the final plot using ggplot2 (Wickham 2016). Here it is possible to alter most of the display options, such as the types of lineages to be shown (observed, hidden, total), as well as their colours and shapes (see e.g. Fig. S8).

**Table S2:**
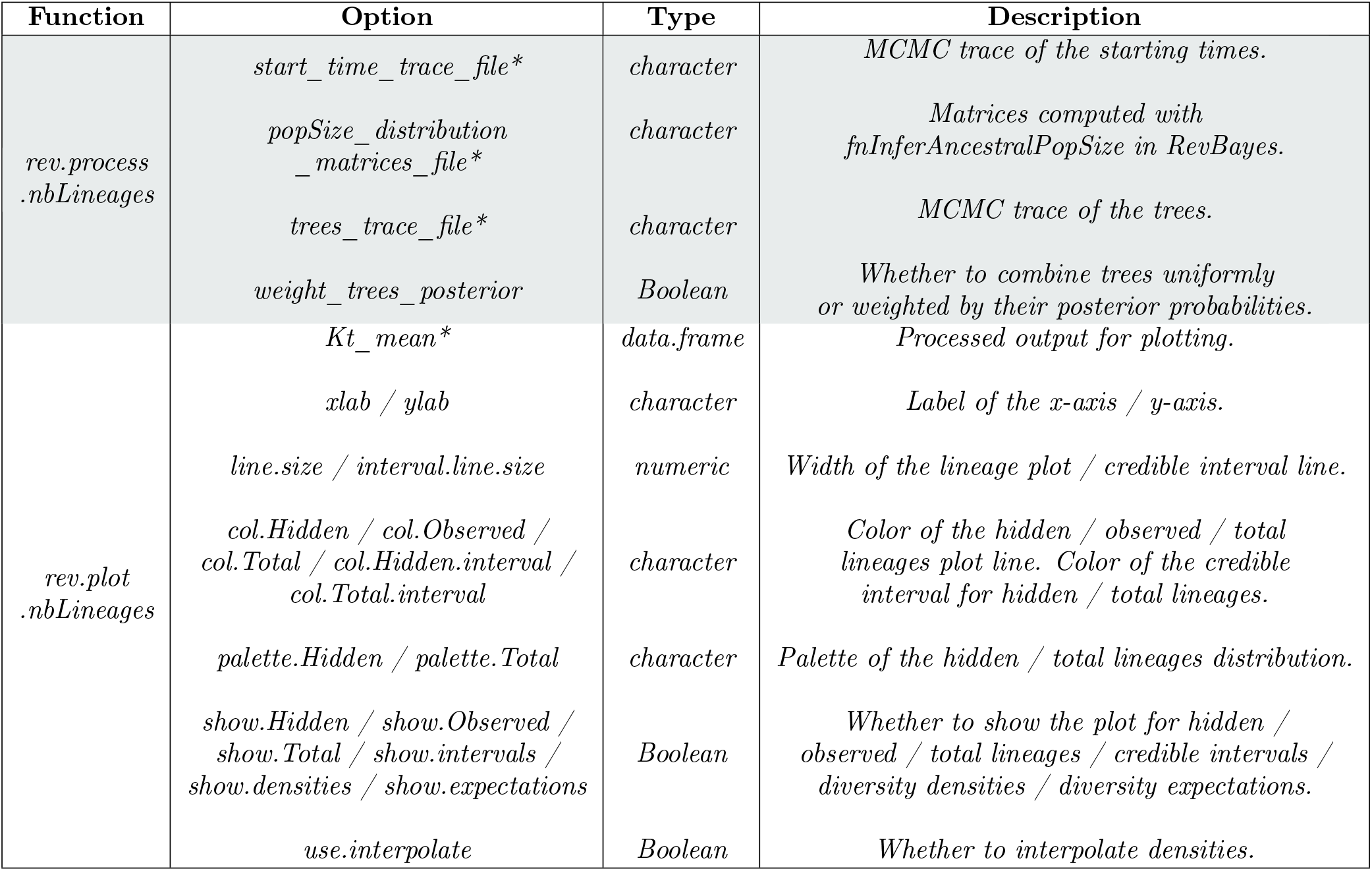
Description of two novel RevGadgets functions for visualizing OBDP diversity-through-time estimations. The input objects and display parameters are detailed, those with an asterisk always have to be provided while the others have default values.

## C Qualitative validation: “blind test” on simulated data

Parameter values used to simulate the two datasets used in the blind test are presented in Table S3. Two trees with occurrences have been simulated under the OBDP (parameters 1-6). For “dataset 1”, genetic sequences along the first tree are simulated according to a K80 model of molecular evolution (parameters 7-9) and recorded only for extant taxa. Binary traits are simulated according to a Markov process with symmetrical rates (parameters 10-12) and are recorded for both extant and extinct taxa. This corresponds to a classic macroevolution scenario. For “dataset 2”, genetic sequences along the second tree are simulated according to a K80 model of molecular evolution (parameters 7-9) and recorded for extant and extinct individuals. This allows us to have a better resolution of the underlying tree than in the first dataset. Moreover, getting genetic sequences for individuals sampled in the past corresponds more to an epidemiology scenario.

**Table S3:** Parameter values used to simulate two datasets and test our OBDP inference workflow.

| Parameter values |  |  |  |  |  |  |  |  |  |  |  |
| --- | --- | --- | --- | --- | --- | --- | --- | --- | --- | --- | --- |
| $\lambda$ | $\mu$ | $\psi$ | $\omega$ | $r$ | $\rho$ | $m_{nt}$ | $\alpha_{nt}$ | $\beta_{nt}$ | $m_{morpho}$ | $q_{01}$ | $q_{10}$ |
| 1 | 0.9 | 0.2 | 0.3 | 0 | 0.8 | 10000 | 0.01 | 0.02 | 60 | 0.03 | 0.03 |

Two of us, ignorant of the priors Joelle: values used for simulation, designed the inference protocol and conducted the analysis, taking as input the occurrences, sequences, and morphological data only. Priors used for inference on “dataset 1” are presented in Table S4 and the general setup for analysis is illustrated in Figure S7. Priors used for inference on “dataset 2” were very similar, except for the absence of a model of morphological evolution, and they are presented in Table S5.

**Figure S7:**
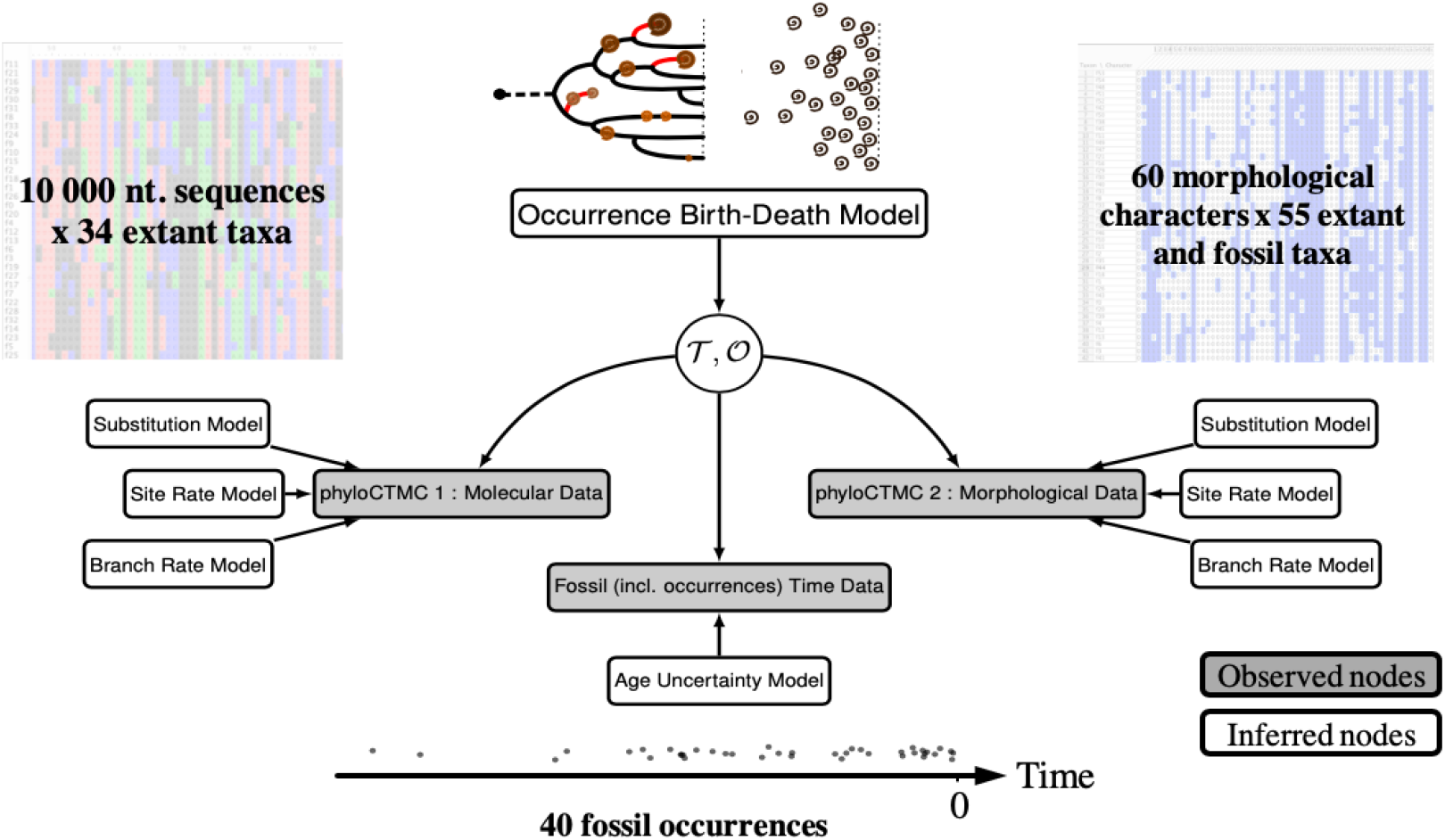
Modular representation of the graphical models used in the qualitative validation analysis. Modified from Heath et al. (2019). The simulated data, noted in the grey nodes are used to deduce the posterior distributions of all other random variables noted in the white nodes.

**Table S4:** Prior distributions on the OBDP parameters and models for the “Blind Test” analysis on dataset 1. Notations: 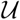 for the Uniform distribution, *Exp* for Exponential, *Dir* for Dirichlet, *GTR* for the General Time Reversible substitution model and *MK* for the Mk model, the analog of JC69 for an arbitrary number of character states.

| Parameter | Prior | Model | Prior |
| --- | --- | --- | --- |
| $\lambda$ | $Exp(10)$ | *Molecular evolution:*<br>$GTR + \Gamma$ | Strict clock rate: $Exp(10)$ |
| $\mu$ | $Exp(10)$ | | Exchangeability rates: $Dir(1, 1, 1, 1, 1, 1)$ |
| $\psi$ | $Exp(10)$ | | Stationary frequencies: $Dir(1, 1, 1, 1)$ |
| $\omega$ | $Exp(5)$ | | Gamma distribution shape: $Exp(1)$ |
| $\rho$ | $\mathcal{U}(0, 1)$ | | |
| $r$ | 0 | Morphological evolution:<br>$MK + \Gamma$ | Strict clock rate: $Exp(1)$ |
| $t_{or}$ | $\mathcal{U}(7.7, 12)$ | | Gamma distribution shape: $Exp(1)$ |

**Table S5:**
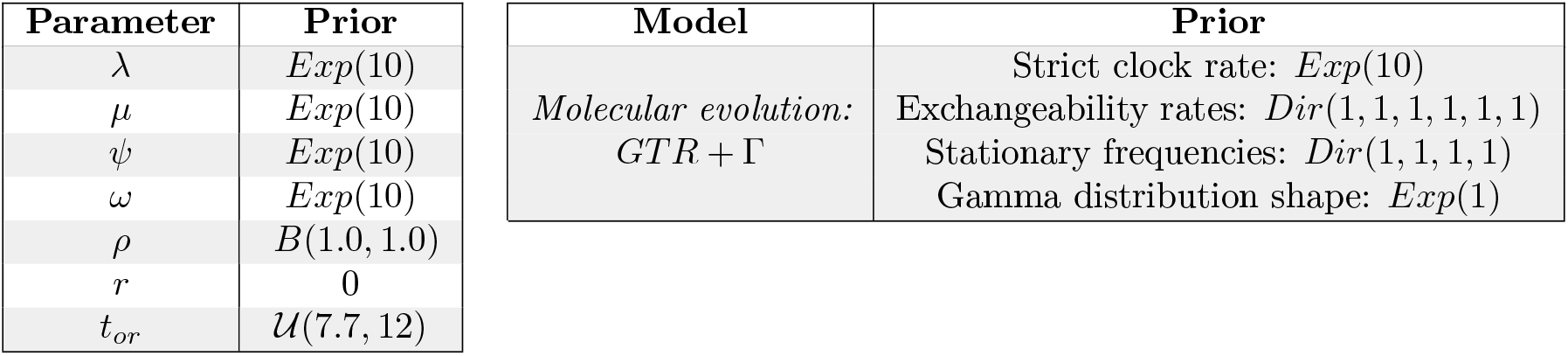
Prior distributions of the OBDP parameters and models for the “Blind Test” analysis on dataset 2. Notations: 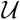 for the Uniform distribution, *B* for the Beta distribution, *Exp* for Exponential, *Dir* for Dirichlet, *GTR* for the General Time Reversible substitution model.

| Parameter | Prior | Model | Prior |
| --- | --- | --- | --- |
| $\lambda$ | $Exp(10)$ | *Molecular evolution:*<br>$GTR + \Gamma$ | Strict clock rate: $Exp(10)$ |
| $\mu$ | $Exp(10)$ | | Exchangeability rates: $Dir(1, 1, 1, 1, 1, 1)$ |
| $\psi$ | $Exp(10)$ | | Stationary frequencies: $Dir(1, 1, 1, 1)$ |
| $\omega$ | $Exp(10)$ | | Gamma distribution shape: $Exp(1)$ |
| $\rho$ | $B(1.0, 1.0)$ | | |
| $r$ | 0 | | |
| $t_{or}$ | $\mathcal{U}(7.7, 12)$ | | |

In our blind inferences, we recovered posterior distribution of diversity trajectories (Fig. S8) and trees (Fig. S9) which are very close to the real data from the simulations. The true number of hidden lineages is most of the time near the expectation of the inferred posterior distribution and more importantly always in the 95% posterior credible interval. When looking at the total number of lineages – i.e. species richness in macroevolution or prevalence in epidemiology – the estimates remains very close to the truth and almost always in the 95% credible interval.

**Figure S8:**
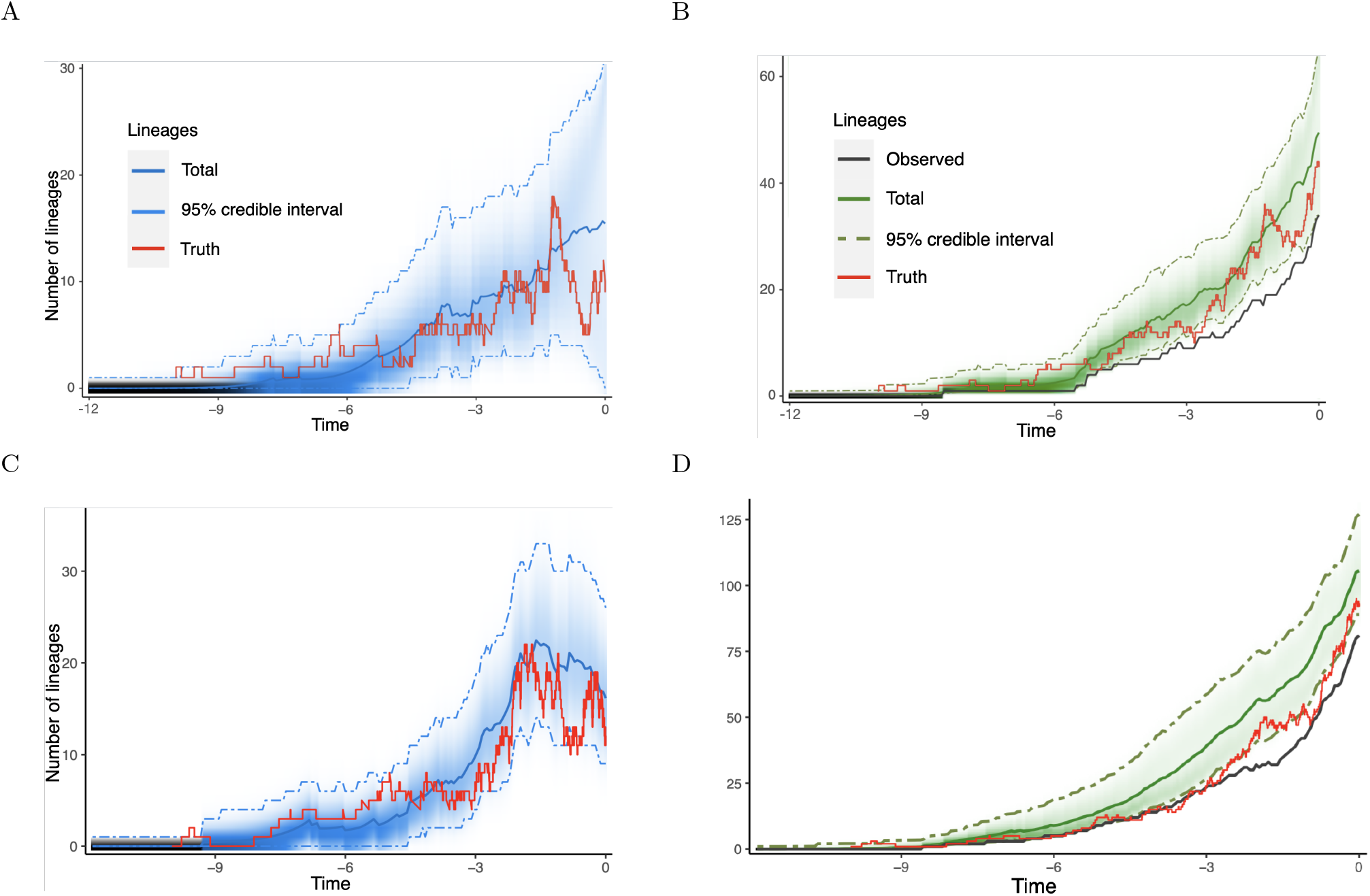
Validation of the diversity dynamics inferred by OBDP compared to the true simulated data. (A) Posterior probability distribution of the number of hidden lineages through time for “dataset 1”, plotted with the new RevGadgets utilities. (B) Posterior probability distribution of the total number of lineages through time for “dataset 1”. (CD) Same as (AB), but for “dataset 2”. The 95% credible intervals are indicated in dashed lines, the expected number of lineages is in blue or green and the true, simulated, trajectory in red. The black line represents the inferred Lineages Through Time (LTT) plot, note that the total diversity equals the LTT plus the hidden diversity.

**Figure S9:**
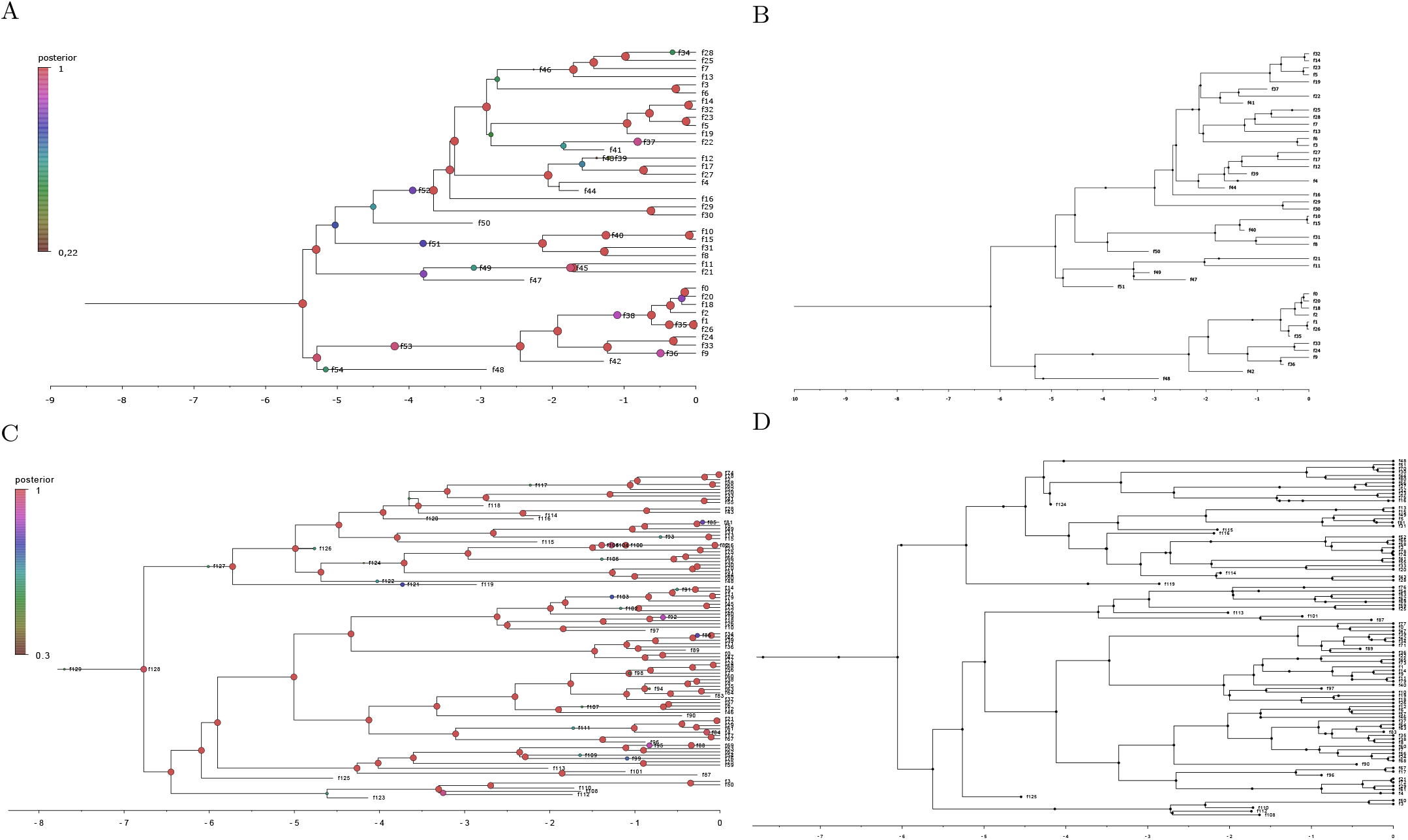
Validation of the inferred trees against the true simulated ones. (A) Inferred phylogenetic tree for “dataset 1”, visualized in FigTree 1.4.4. The node colors refer to their posterior probability. (B) Original simulated tree for “dataset 1”, aligned on the same temporal scale. Note that the topology is well recovered but divergence dates do not always perfectly match. (CD) Same as (AB) but on “dataset 2”. Due to a greater amount of data in genetic sequences of both past and extant individuals, the divergence dates tend to be better inferred.

## D Macroevolution application: Inferring past cetacean diversity

### D.1 Preliminary analysis of the cetacean occurrence fossil record

A detailed notebook is available at https://github.com/Jeremy-Andreoletti/Cetacea_PBDB_Occurrences to follow our exploration of the cetacean dataset. We identified several biases in their fossil record, in particular much more variable occurrence densities – defined as the number of occurrences by unit of time in the stratigraphic range of a clade – than expected from our model (see Figure S10).

Since OBDP assumes that only one individual of a species will be sampled at a time, we subsampled the dataset to aggregate all occurrences of the same taxon found in the same geological formation. This subsampling also reduced the observed discrepancy in occurrence densities. The final subsampled dataset was composed of 968 occurrences.

**Figure S10:**
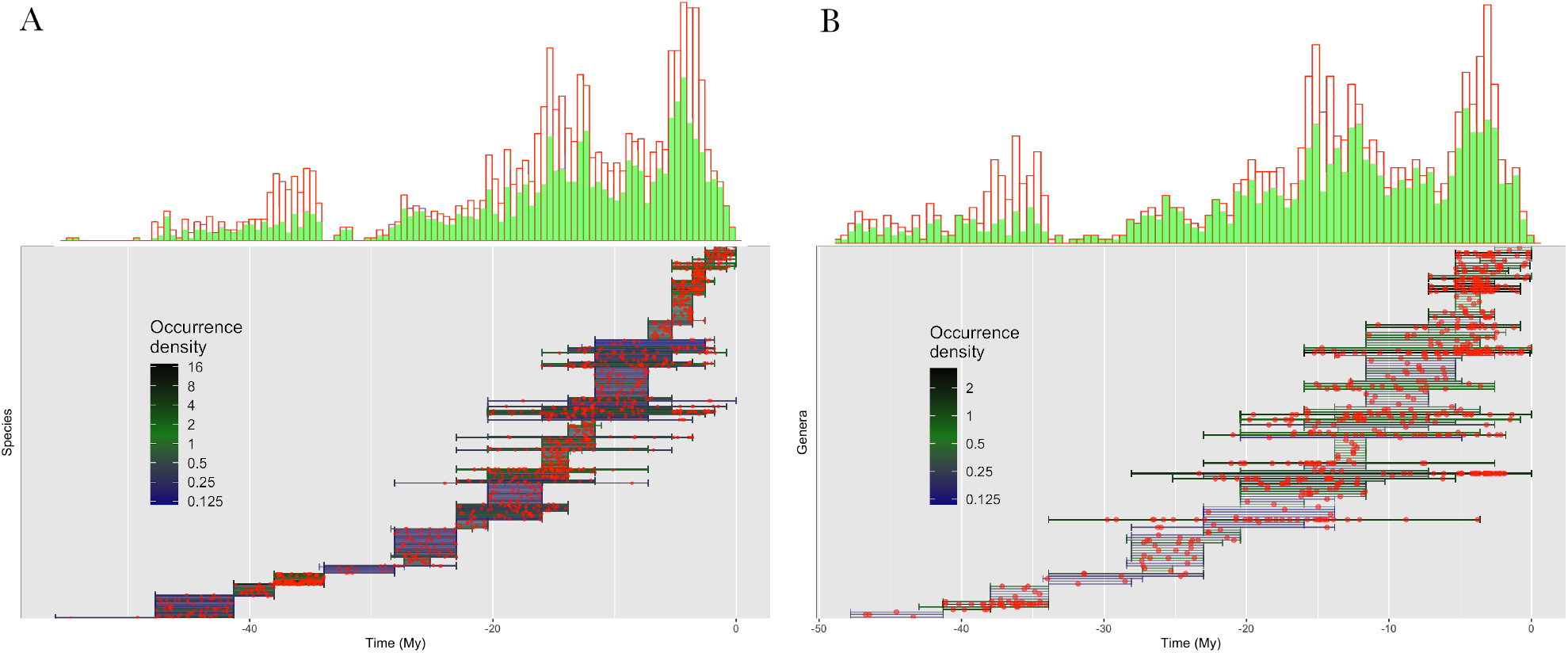
Occurrence distributions and bias correction, for cetacean species (A) and genera (B). At the top, occurrence distributions are compared before (red) and after (green) aggregating in geological formations. Below, stratigraphic ranges are displayed over time and colored according to the density of occurrences (red dots).

### D.2 Detailed priors used for Bayesian inference

We detail in Table S6 all priors used for the inference on the cetacean dataset.

**Table S6:**
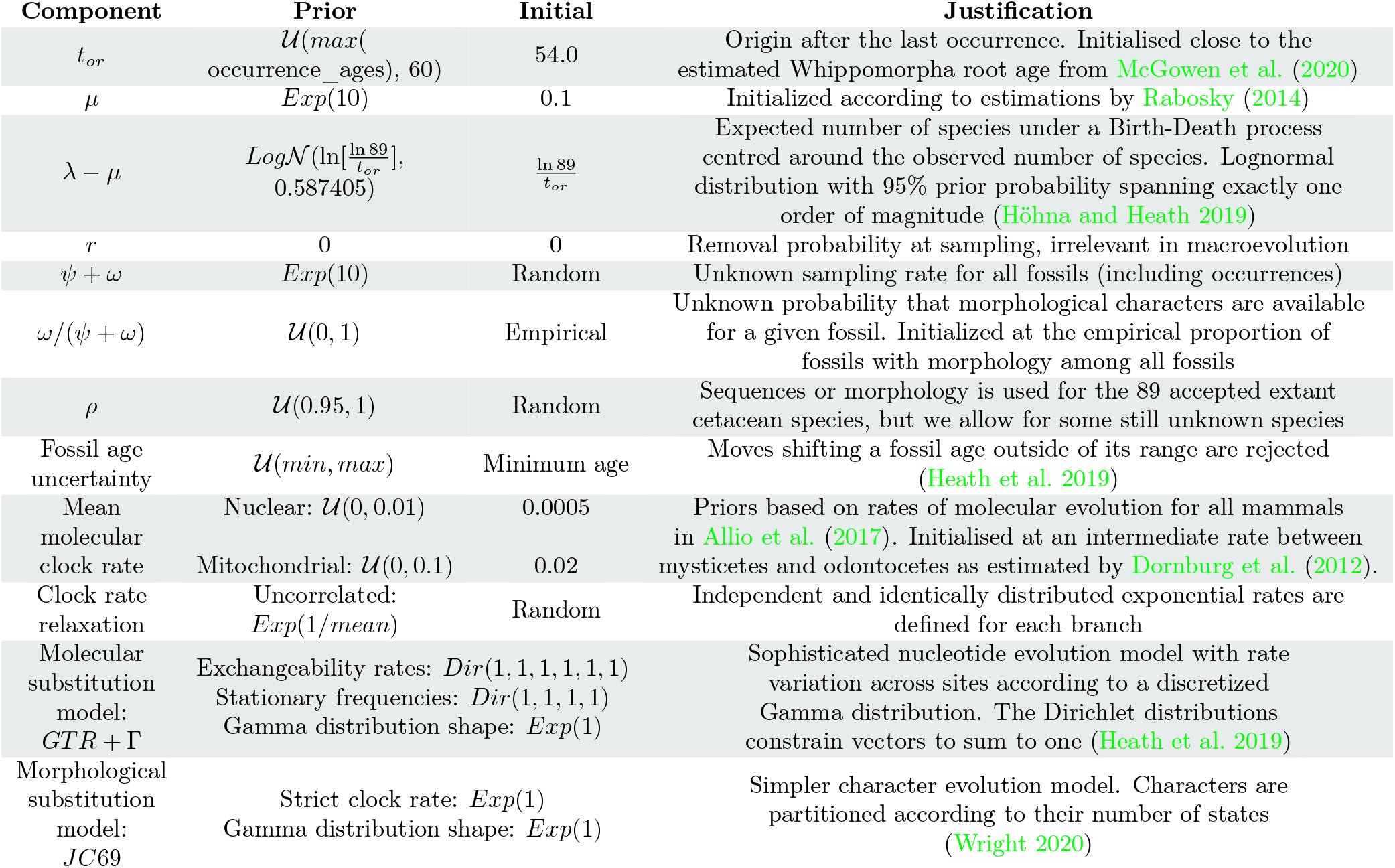
Prior distributions for parameters and models of the Cetacea analysis. For each parameter its prior distribution, its initial value at the origin of the MCMC chain (set to speed up convergence) and the references that support these choices are indicated. Notations: 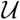 for the Uniform distribution, *Exp* for Exponential, 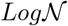 for Log-Normal, *Dir* for Dirichlet, *GTR* for General Time Reversible and *JC*69 for the Jukes-Cantor 1969.

## E Epidemiology application: the Diamond Princess SARS-2 COVID-19 outbreak dynamics

### E.1 Data acquisition on GISAID

We gratefully acknowledge the following Authors from the Originating laboratories responsible for obtaining the specimens, as well as the Submitting laboratories where the genome data were generated and shared via GISAID, on which this research is based. All Submitters of data may be contacted directly via www.gisaid.org

**accession ID** EPI_ISL_416565, EPI_ISL_416566, EPI_ISL_416567, EPI_ISL_416568, EPI_ISL_416569, EPI_ISL_416570, EPI_ISL_416571, EPI_ISL_416572, EPI_ISL_416573, EPI_ISL_416574, EPI_ISL_416575, EPI_ISL_416576, EPI_ISL_416577, EPI_ISL_416578, EPI_ISL_416579, EPI_ISL_416580, EPI_ISL_416581, EPI_ISL_416582, EPI_ISL_416583, EPI_ISL_416584, EPI_ISL_416585, EPI_ISL_416586, EPI_ISL_416587, EPI_ISL_416588, EPI_ISL_416589, EPI_ISL_416590, EPI_ISL_416591, EPI_ISL_416592, EPI_ISL_416593, EPI_ISL_416594, EPI_ISL_416595, EPI_ISL_416596, EPI_ISL_416597, EPI_ISL_416598, EPI_ISL_416599, EPI_ISL_416600, EPI_ISL_416601, EPI_ISL_416602, EPI_ISL_416603, EPI_ISL_416604, EPI_ISL_416605, EPI_ISL_416606, EPI_ISL_416607, EPI_ISL_416608, EPI_ISL_416609, EPI_ISL_416610, EPI_ISL_416611, EPI_ISL_416612, EPI_ISL_416613, EPI_ISL_416614, EPI_ISL_416615, EPI_ISL_416616, EPI_ISL_416617, EPI_ISL_416618, EPI_ISL_416619, EPI_ISL_416620, EPI_ISL_416621, EPI_ISL_416622, EPI_ISL_416623, EPI_ISL_416624, EPI_ISL_416625, EPI_ISL_416626, EPI_ISL_416627, EPI_ISL_416628, EPI_ISL_416629, EPI_ISL_416630, EPI_ISL_416631, EPI_ISL_416632, EPI_ISL_416633, EPI_ISL_416634, EPI_ISL_454749
**Originating Laboratory** Japanese Quarantine Stations
**Submitting Laboratory** Pathogen Genomics Center, National Institute of Infectious Diseases
**Authors** Tsuyoshi Sekizuka, Kentaro Itokawa, Rina Tanaka, Masanori Hashino, Tsutomu Kageyama, Shinji Saito, Ikuyo Takayama, Hideki Hasegawa, Takuri Takahashi, Hajime Kamiya, Takuya Yamagishi, Motoi Suzuki, Takaji Wakita, Makoto Kuroda

**Figure S11:**
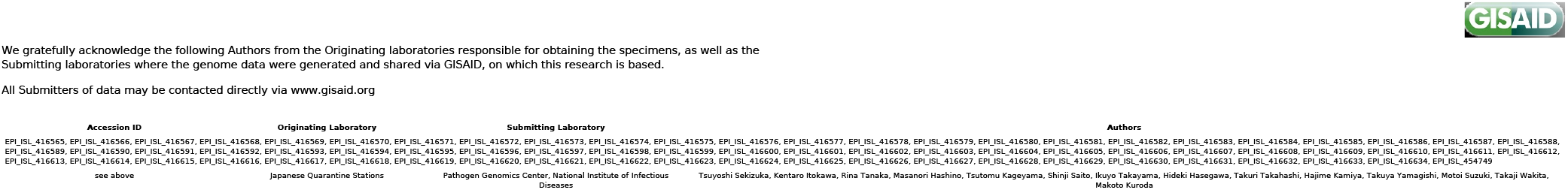
Genome sequences used, originating and submitting labs generated on GISAID. Content is reproduced above.

### E.2 Pre-processing the data

All case count and sequencing data were available at a resolution of days.

In order to use the main method described in this article, the case count record had to be pre-processed so that occurrences are spread throughout the days. For a day with a case count of *n* newly infected individuals, we drew *n* time points uniformly distributed throughout the day. The resulting dataset is shown in Figure S12.

**Figure S12:**
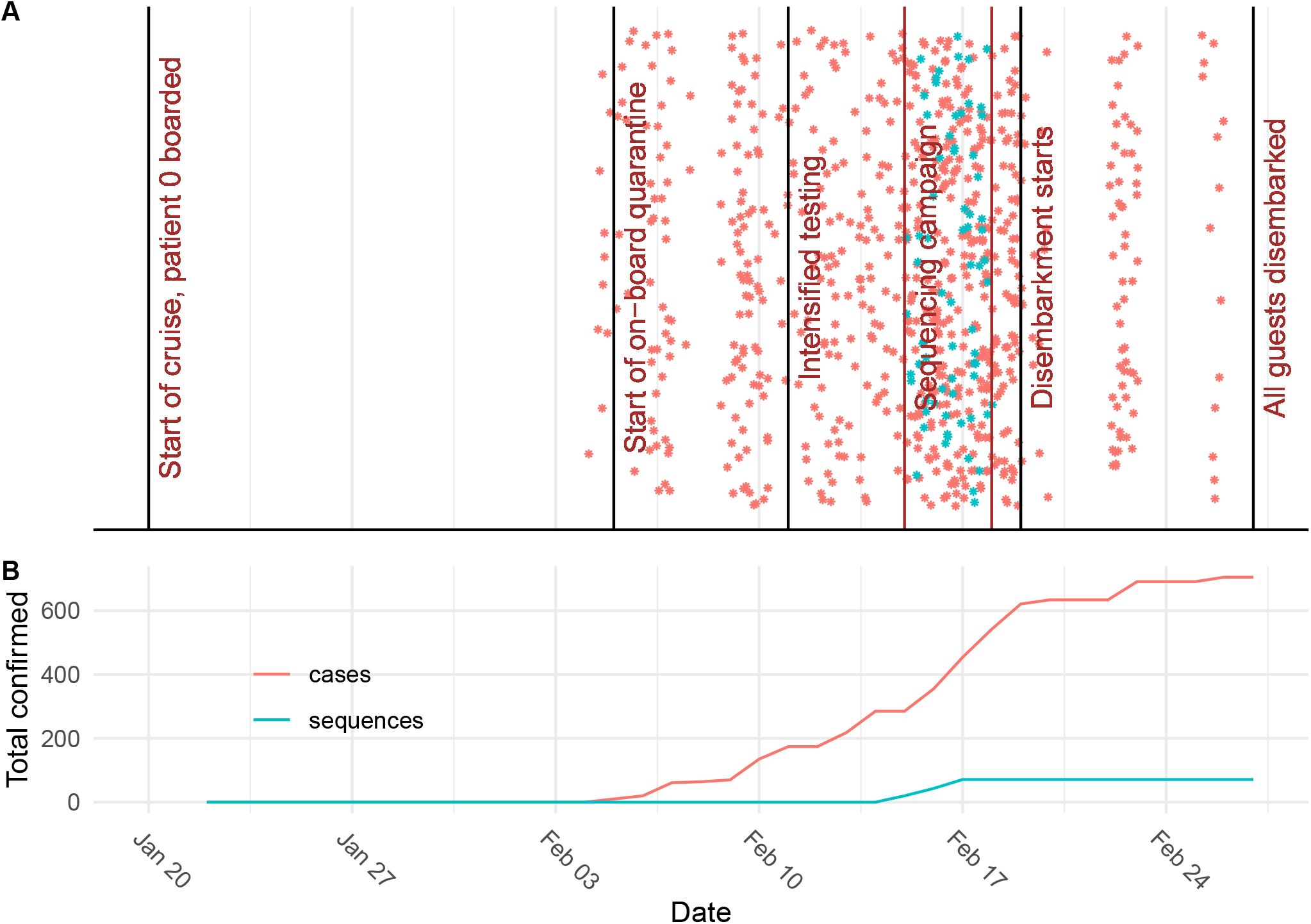
Pre-processed dataset for the Diamond Princess outbreak analysis. (A) Exact dates assigned to occurrences and sequences for the analysis. (B) Total case counts and sequences through time.

**Figure S13:**
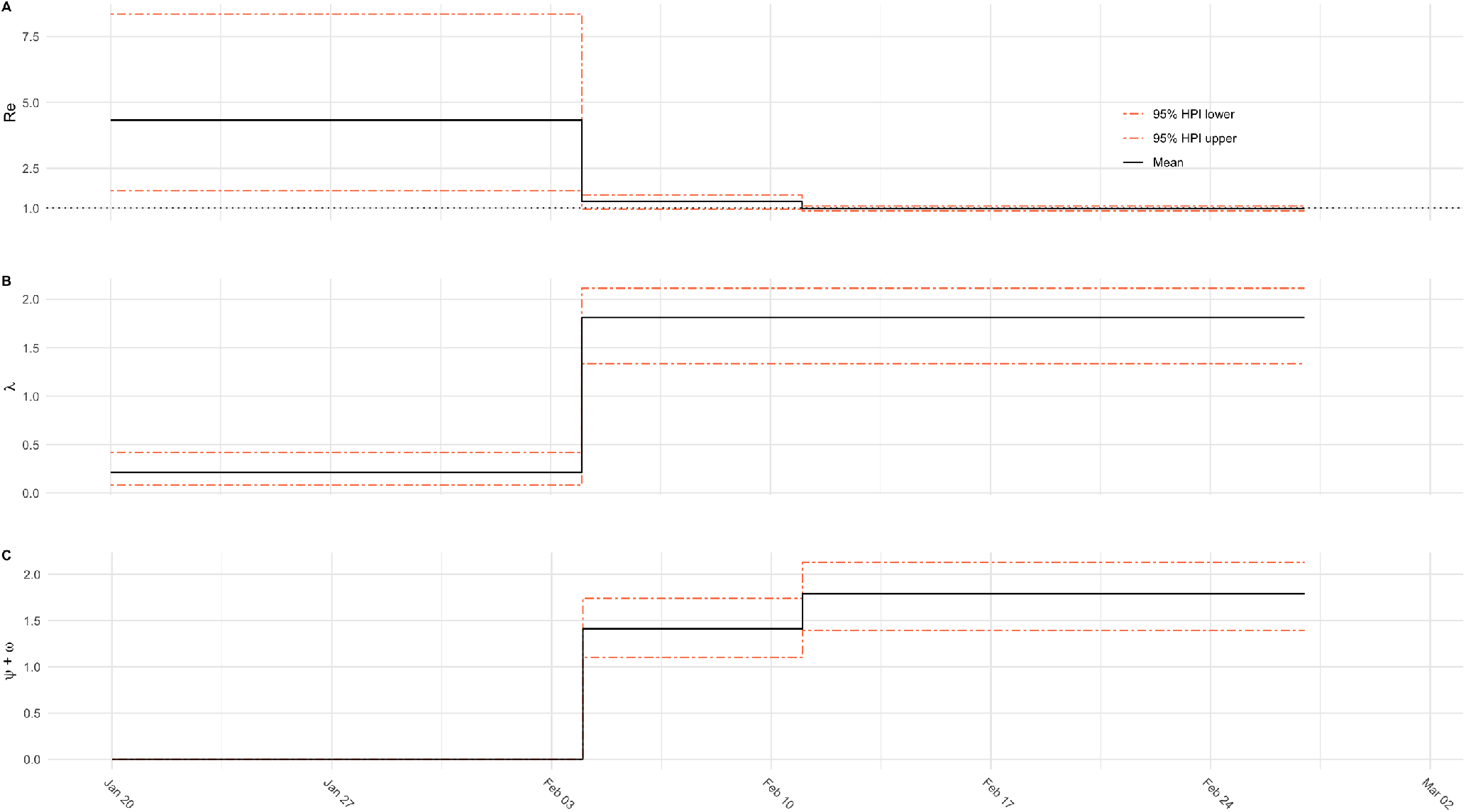
Detailed parameter estimates obtained from the COVID-19 outbreak analysis. (A) Reproductive number estimates. (B) Birth rate estimates. (C) Total sampling (sequencing and PCR testing) rate estimates.

### E.3 Detailed priors

We detail in Table S7 all priors used for the inference on the outbreak dataset of COVID-19 aboard the Diamond Princess.

The mean of the prior distribution of *ψ* + *ω* is set up to be the number of tests used on the ship, per day and per passenger, on the two periods.

- Within the first 7 days period, from February 4th to February 11th, there were 439 tests carried out, on 3711 passengers, leading to 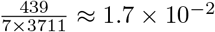 tests per day per passenger.
- on the following 15 days period, from February 11th to February 27th, there were 3622 tests carried out, on 3711 passengers, leading to 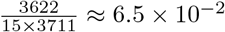 tests per day per passenger.

**Table S7:**
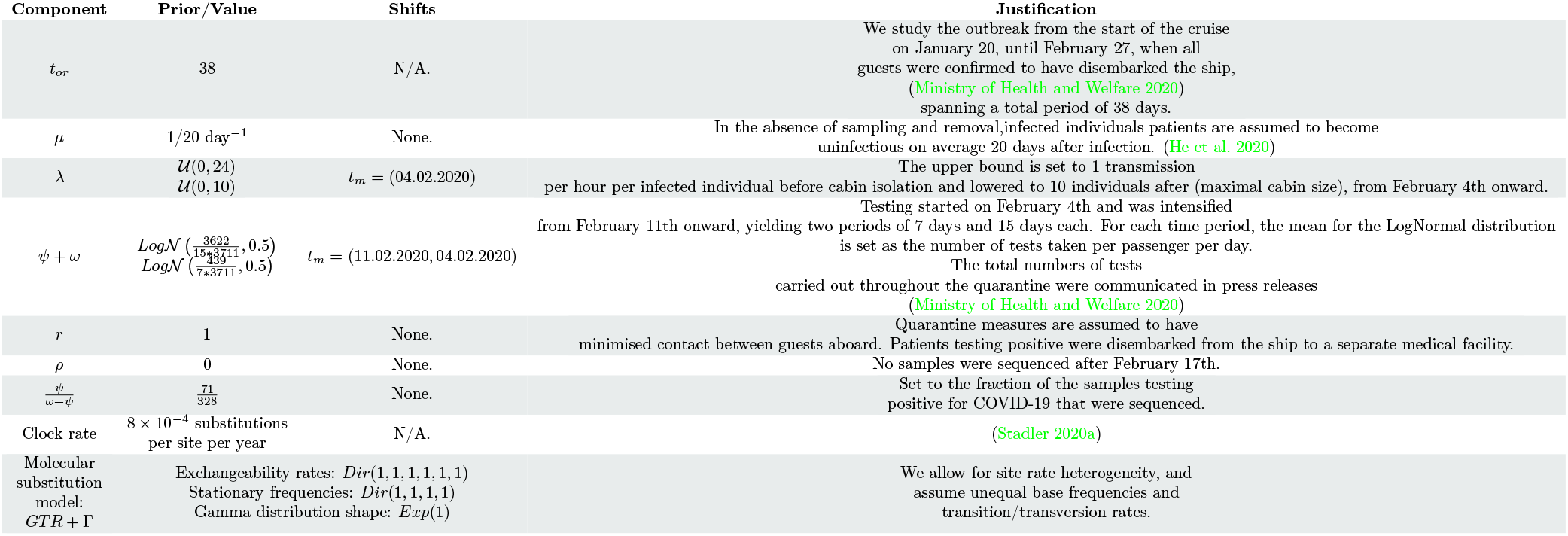
Prior distributions for parameters and models of the SARS-2 COVID-19 analysis. For each parameter its prior distribution or value and the references that support these choices are indicated.

